# Modulation of the GT Family 47 clade B gene affects arabinan deposition in elaters of *Marchantia polymorpha*

**DOI:** 10.1101/2025.05.02.651808

**Authors:** H. S. Kang, E. R Lampugnani, X. Tong, P.K. Prabhakar, E. Flores-Sandoval, J. Hansen, B. Jørgensen, J.L. Bowman, B.R. Urbanowicz, B. Ebert, S. Persson

## Abstract

The plant cell wall polymer α-1,5-linked arabinan is associated with important functions in plant physiology, such as cell wall flexibility and desiccation tolerance in angiosperms. However, its biosynthetic mechanism remains obscure. A putative arabinan arabinosyltransferase (AraT) is the Arabidopsis ARABINAN-DEFICIENT1 in GT family 47 clade B (GT47B), but its role is inferred by its mutant cell wall phenotypes, and not through *in vitro* assays. With seven other genes in the clade, investigating the function of the members in GT47B is hampered by the high genetic redundancy in Arabidopsis. Instead, we probe the function of the only two genes in GT47B in the liverwort model organism *Marchantia polymorpha,* named *MpARAD-Like 1* (*MpARADL1*) and *MpARADL2*. *Mparadl1* and *Mparadl2* loss-of-function mutants and overexpression lines were generated, and their cell wall composition was probed using Comprehensive Microarray Polymer Profiling (CoMPP), glycosyl linkage analysis and immunolabelling. Interestingly, *Mparadl2* mutants have much less of the 1,5-α-L-arabinan epitope in elaters since LM6 antibody recognition showed a notable reduction. However, 5-linked Ara*f* levels in the Marchantia thallus tested with glycosyl linkage analysis are comparable between mutants and wild type, suggesting that there is another enzyme forming 5-Ara*f* linkages in *Marchantia.* Our attempts to obtain the biochemical activity of these enzymes through the expression and purification of AtARAD1, MpARADL1 and MpARADL2 proteins in the HEK293 cell heterologous expression system were unsuccessful. Therefore, we suspect that an evolutionarily conserved, and clade-specific mechanism is required to confer solubility for purification and subsequent activity characterisation of GT47B proteins. Previous studies have shown that co-expression of protein interaction partners can enhance protein solubility in the HEK293 cell system, but co-expression of MpARADL2 with AtARAD1 and MpARADL1 did not yield soluble proteins, prompting further investigation into the protein-protein interactors of GT47B proteins.

## Introduction

The plant cell wall is a rigid yet extensible extracellular layer rich in polysaccharides. Plant cell wall polysaccharides broadly group into cellulose, which forms microfibrils that act as scaffolding, and hemicelluloses, pectins and glycoproteins which form the hydrated matrix of the cell wall where the cellulose microfibrils are embedded (Lampugnani et al. 2018). Arabinan is a cell wall polymer typically associated with the pectic polysaccharide rhamnogalacturonan-I (RG-I), one of three major domains of pectin with homogalacturonan and RG-II, and arabinogalactan proteins (AGPs), which are major structural glycoproteins found in plant cell walls (Wachananawat et al. 2020; Silva et al. 2020). Arabinan consists of a linear α-1,5-L-arabino*furanose* backbone that can be branched at the *O*-2, *O*-3 and/or *O*-5 position (Cardoso et al., 2002, 2007; Dourado et al. 2006; Wefers et al., 2014, 2018). In angiosperms, the *in vivo* functions of arabinan encompass many aspects of plant physiology (Cankar et al. 2014; Carroll et al. 2022; Jones et al. 2003; Peña and Carpita 2004; Verhertbruggen et al. 2013) including mechanical stress tolerance in desiccation tolerant plants (Moore et al. 2013; Moore, Farrant, and Driouich 2008). To better understand the diverse roles of cell wall arabinan and unravel the relationship between its structure and function, it is necessary to elucidate its biosynthetic mechanism.

Plant cell wall polysaccharides, including arabinan, are built by a class of enzymes called glycosyltransferases (GTs), which utilise nucleotide sugar donors to glycosylate specific acceptor moieties (Lairson et al. 2008). GTs are catalogued in an online database (CAZy; Cantarel et al. (2009); Lombard et al. (2014)) and classified into over 130 different families. Among them, cell wall biosynthetic GTs have been characterised in 15 different families, (Mariette et al. 2023) including GT family 47 (GT47), where several cell wall-related GTs have been identified (Zhang et al. 2023). GT47 is divided into six clades (A-F), including clade B, which harbours a putative α-1,5-arabinan arabinosyltransferase (AraT) called At *ARABINAN DEFICIENT 1* (AtARAD1) (Harholt et al. 2006). The *Arabidopsis* T-DNA insertional mutant *Atarad1* showed reduced levels of 5-linked Ara*f* through the LM6 α-1,5-arabinan antibody recognition and cell wall compositional analysis of mutant stem sections. The authors further showed that its homolog, AtARAD2 functions in branching sugars onto arabinan in an AtARAD1 dependent manner (Harholt et al., 2012). Furthermore, Carroll et al., (2022) has shown that the overexpression of AtARAD1 leads to increased LM6 binding in the stomata, and Lampugnani and colleagues (2016) have shown that Arabidopsis plants heterologously overexpressing the *Nicotiana alata* ARAD-Like 1 (*NaARADL1*) produce arabinan-rich root exudates. However, the biochemical function of these putative arabinan AraTs have not been demonstrated, and their role in arabinan biosynthesis remains circumstantial.

To tackle this gap in our knowledge, we employed a reverse genetics approach using the liverwort model organism, *Marchantia polymorpha* (Figure 1). The low genetic redundancy of *Marchantia* is being increasingly appreciated for unravelling the complexity of cell wall biosynthesis (Honkanen et al. 2016; Wachananawat et al. 2020). While Arabidopsis has eight members in GT47B, Marchantia has only two orthologs, named MpARAD-Like (MpARADL1; Mp5g02380) and MpARADL2 (Mp3g05600). In this study, we investigate the *in vivo* function of Marchantia GT47B proteins and attempt to purify members of GT47B in a novel heterologous expression system to advance towards obtaining soluble proteins suitable for *in vitro* catalytic assays.

**Figure 1.**
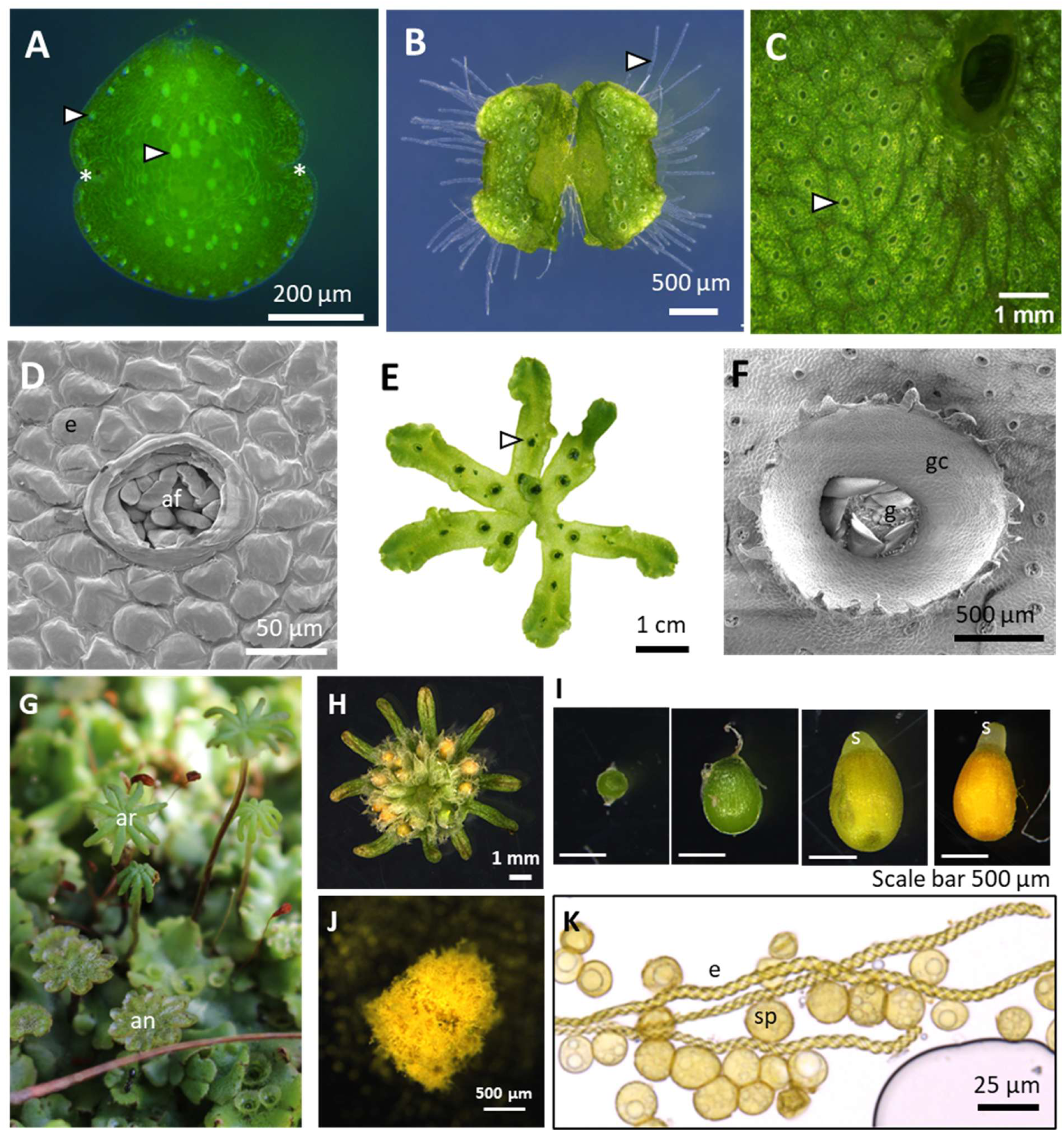
Morphology of *Marchantia polymorpha*. A) Dormant gemmae, a clonal structure of *M. polymorpha*. Arrowheads = oil bodies, asterisk = apical notch (meristem). B) 7d old *M. polymorpha* gemmae. Arrowhead = rhizoids. C) Air pores (arrowhead) are openings to air chambers, where gas exchange occurs. D) Scanning electron microscope (SEM) image of an air pore. e = epidermal cell, af = assimilatory filaments. E) 4-week old *M. polymorpha* thallus. Gemmae cups form along the thallus (arrowhead). F) SEM image of a gemmae cup (gc). g = gemmae. G) The reproductive structures of *M. polymorpha*. ar = archegoniophore (female), an = antheridiophore (male). H) Fertilised archegoniophore with mature spore capsules (sporophytic life stage of *M. polymorpha*). I) Progression of spore capsule maturation. S = seta, the foot of the spore capsule which extends upon maturation. J) Mature spore capsules burst and assisted by elaters, disperse spores. (K) Spores (Sp) and elater cells (e) of *M. polymorpha*.

## Results and Discussion

### Identification of MpARADL1 and 2 and characterisation of their tissue-wide and subcellular expression patterns

Orthofinder analysis across nine species of the Viridiplantae clade identified two Marchantia orthologs of the Arabidopsis GT47B genes (Kang et al., 2025), Mp5g02380 (MpARADL1) and Mp3g05600 (MpARADL2). Evolutionary analysis by the Maximum Likelihood method indicates that AtARAD1 and 2 are not the closest orthologs of MpARADL1 or 2 in Arabidopsis (Figure 2). The expression levels of *MpARADL1* and *2* across different tissue types in Marchantia were analysed using quantitative Reverse Transcription Polymerase Chain Reaction (qRT-PCR) (Figure 3). Relative to the thallus, *MpARADL1* showed increased expression in the stalk, gemmae, apical notch, archegoniophore, sporelings and antheridiophore (Figure 3A). *MpARADL2* expression was higher in the stalk, gemmae, sporophyte and antheridiophore when expression levels were normalised to the thallus (Figure 3B). As arabinan is abundant in the sporophytes of Marchantia (Kang et al., 2025), the expression levels of *MpARADL1* and *2* in the developing sporophytes were also analysed with qRT-PCR (Figure 3B). *MpARADL2* expression was higher relative to the thallus at both the developing sporophyte stage and the mature sporophyte stage. Further probing the expression levels in the thallus with the transcriptional reporter lines encoding either the mVENUS yellow fluorescent protein (YFP) or the β-glucuronidase (GUS) gene indicated that *MpARADL1* and *2* were expressed in the apical notch, rhizoids and sperm (Figure 4). *MpARADL2* expression in day 0 gemmae also showed expression in oil body cells (Figure 4A). As putative arabinan GTs, MpARADL1 and 2 are expected to localise to the Golgi apparatus. The coding sequences of MpARADL1 and 2 were C-terminally fused to the mNEON green fluorescent protein (GFP) and co-expressed with the constitutively expressed Golgi marker, which consists of the transmembrane domain of the soybean mannosidase I (ManI) C-terminally fused with RFP (Figure 5). The signal from the translationally fused MpARADL1 and 2 overlapped with the punctate pattern of the Golgi marker when imaged in the dormant gemmae of stably transformed Marchantia plants.

**Figure 2.**
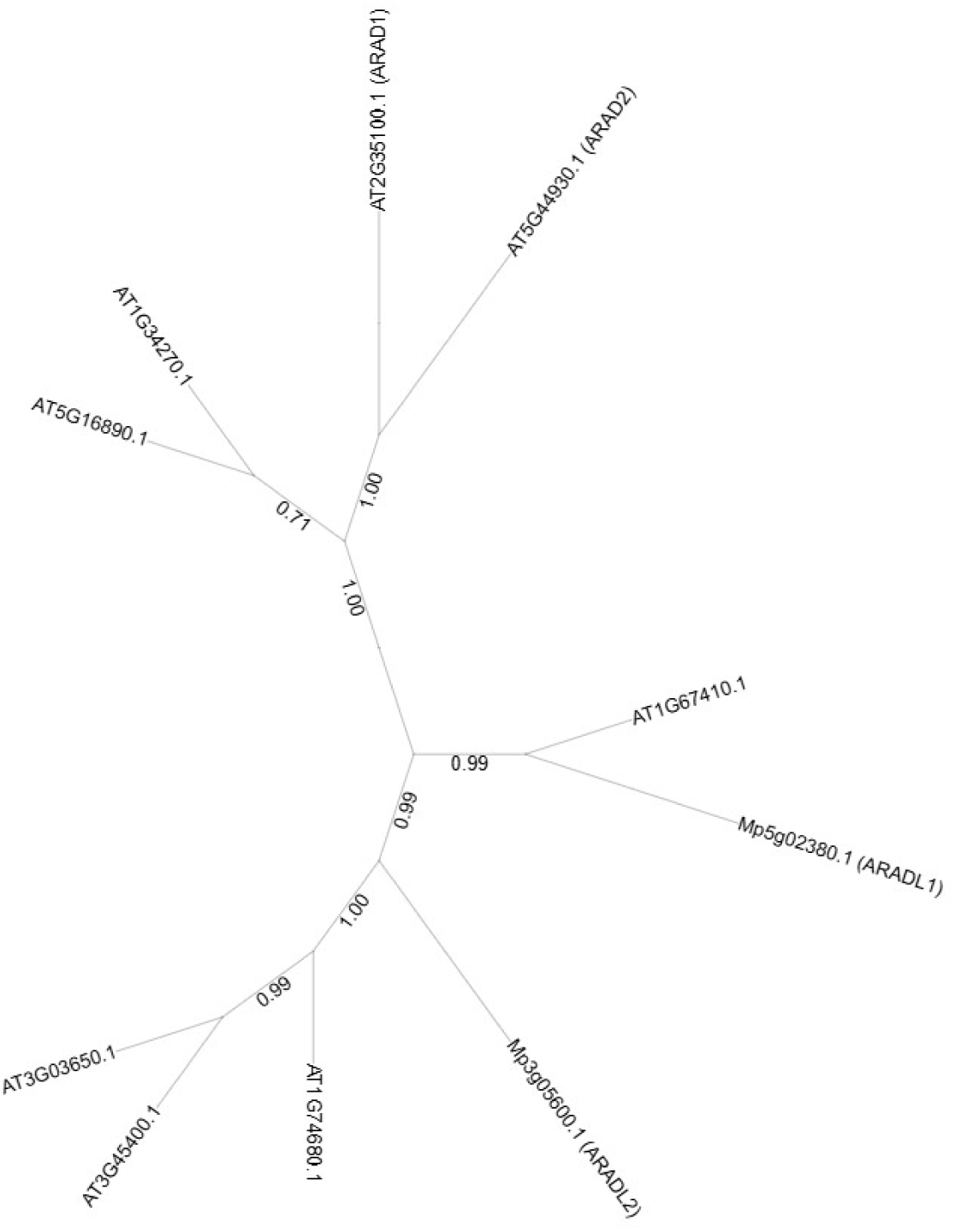
Phylogenetic tree of Arabidopsis and Marchantia GT47B proteins. The evolutionary history was inferred by using the Maximum Likelihood method and Whelan and Goldman model (Whelan and Goldman 2001). The tree with the highest log likelihood (-9693.45) is shown. Initial tree(s) for the heuristic search were obtained automatically by applying Neighbor-Join and BioNJ algorithms to a matrix of pairwise distances estimated using the JTT model, and then selecting the topology with superior log likelihood value. Evolutionary analyses were conducted in MEGA X (Kumar et al., 2018).

**Figure 3.**
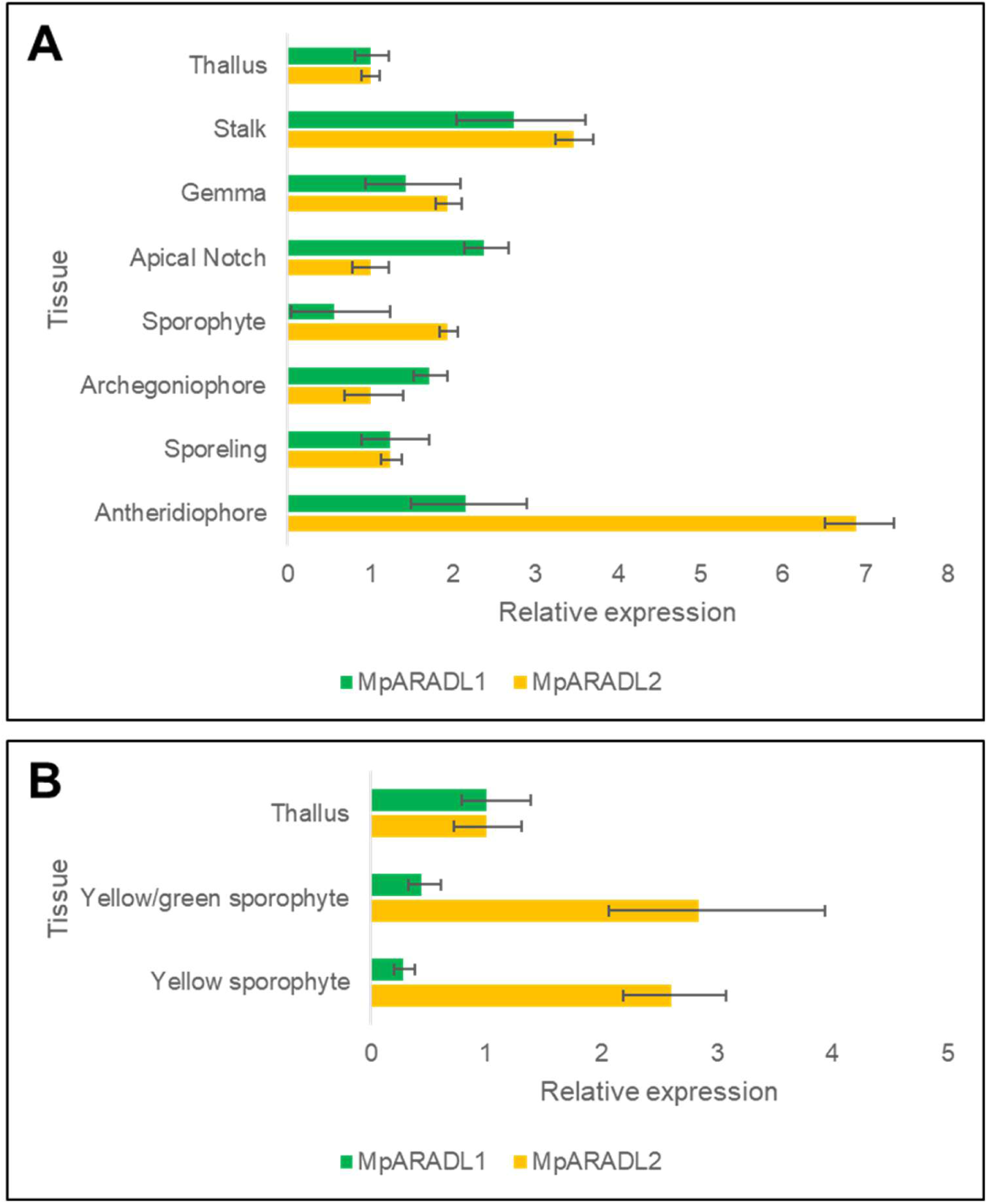
Measurement of relative expression of *MpARADL1* and *2* using quantitative Reverse Transcription Polymerase Chain Reaction (qRT-PCR) across different tissue types of Marchantia.

**Figure 4.**
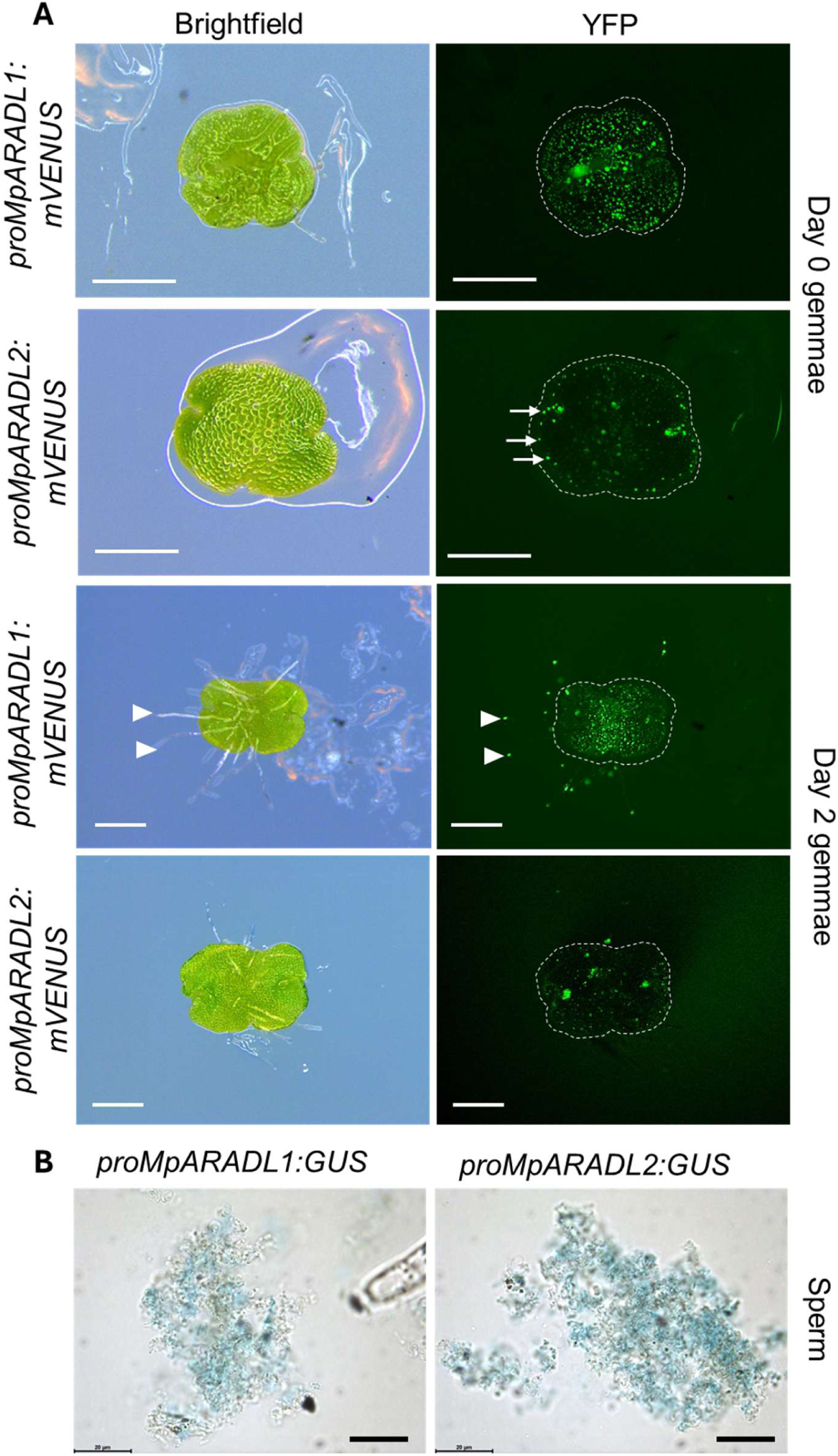
Native promoter constructs of *MpARADLs* driving the expression of either the (A) mVENUS or the (B) β-glucuronidase (GUS) gene. Scale bars (A) = 500 µm, (B) = 20 µm.

**Figure 5.**
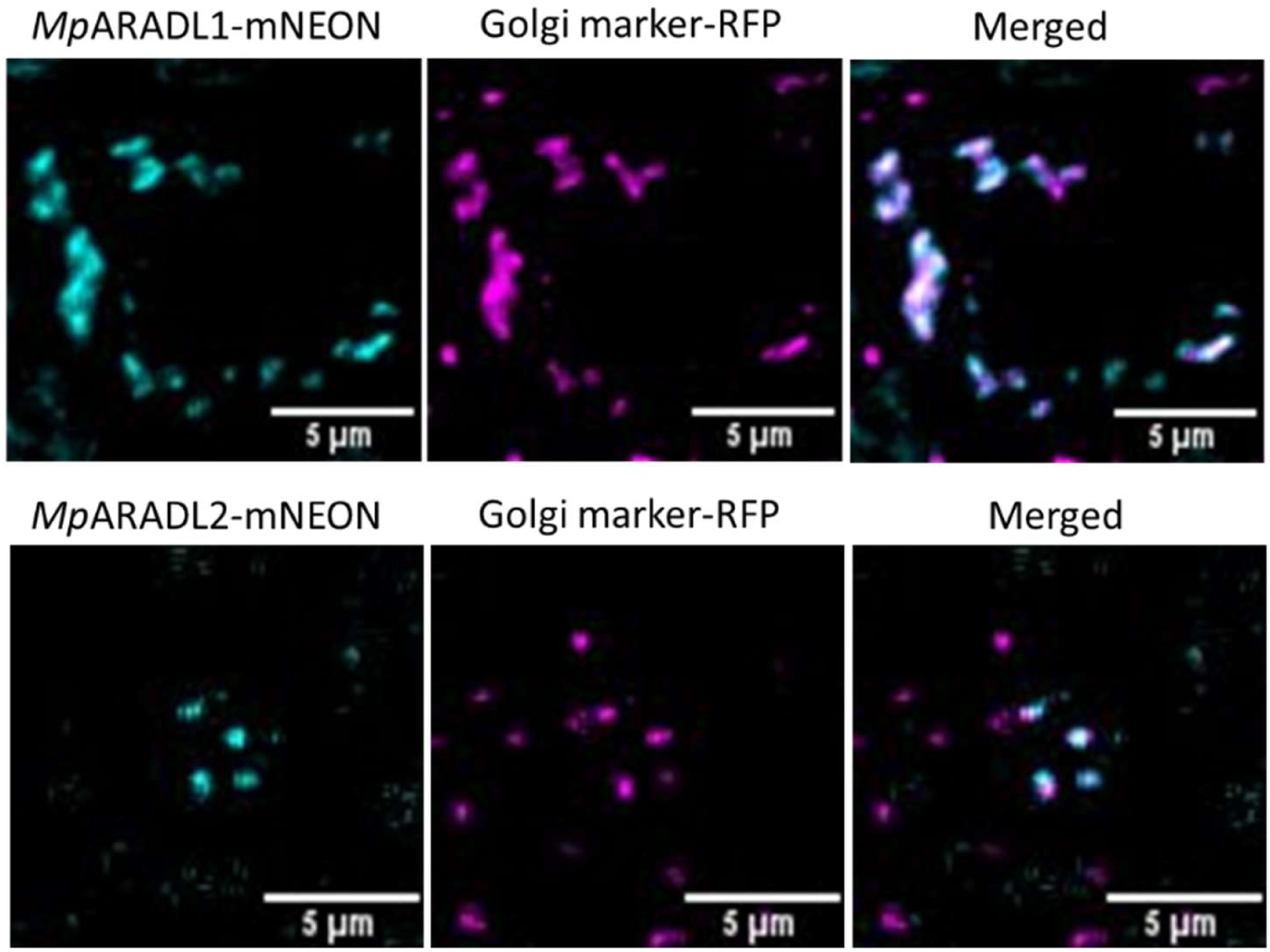
Images of native promoter expressed MpARADL1 and 2, C-terminally fused to mNEON (GFP), co-expressed with the constitutively expressed soybean mannosidase I (ManI) transmembrane domain that is C-terminally fused to mCHERRY (RFP) as the Golgi marker. Images were taken in dormant gemmae of stably transformed Marchantia lines.

### MpARADL1 and 2 play non-redundant roles in plant development

To investigate the function of MpARADL1 and 2 *in planta*, loss-of-function mutants, *Mparadl1* and *Mparadl2* were generated using CRISPR (Clustered Regularly Interspaced Short Palindromic Repeats) mediated gene editing (Figure 6A). Sequencing of the genomic DNA showed that *Mparadl1* has a 53bp insertion in the guide RNA (gRNA) target region, which generated a premature stop codon at AA61. Sequencing of the *Mparadl2* genomic DNA showed a 1bp deletion in the gRNA target region, which led to a premature stop codon at AA104. Overexpression (OE) lines of *MpARADL1* and *2* were generated to understand the gain-of-function effects of *MpARADL1* and *2*, and their overexpression at a transcript level was confirmed with qRT-PCR (Figure 6B). Under normal growth conditions, *Mparadl2* was slightly smaller than wild type (Figure 7), and the *MpARADL1*-OE lines consistently exhibited a phenotype where the air chambers were less compactly organised than wild type (Figure 8). To investigate the clade-wide function of GT47B *in planta*, we employed artificial microRNAs (amiRNA) targeting both MpARADL1 and 2 to induce a simultaneous downregulation of both genes (Figure 9A). amiRNA expressing transformants that were analysed in this study were from the T1 generation, as some transformed sporelings did not develop gemmae. When bulking tissue material for experiments, amiRNA expressing transformants were continuously re-plated on growth media containing fresh antibiotics to suppress the growth of untransformed cells. Some amiRNA induced transgenic lines showed a severe growth phenotype. When the mutants displaying the most severe phenotypes were investigated with scanning electron microscopy (SEM), malformed thalli and ectopic growth of rhizoids was apparent (Figure 9C). The severity of the phenotype appeared correlate with the expression levels of *MpARADL1* (Figure 9A, B). To investigate whether the phenotypes were specific to the reduced expression of *MpARADL1* or *2*, native promoter constructs carrying amiRNA resistant copies of *MpARADL1* and *2* were co-transformed with the amiRNA construct (Figure 9D). The resulting T1 transformants were classified according to thallus morphology. When the amiRNA resistant construct was co-transformed with the constructs carrying the resistant copies of *MpARADL1* or *2*, the proportion of T1 transformants displaying class IV phenotypes was reduced, indicating that the downregulation of MpARADL1 and 2 is likely to be responsible for the phenotype of the amiRNA mutants.

**Figure 6.**
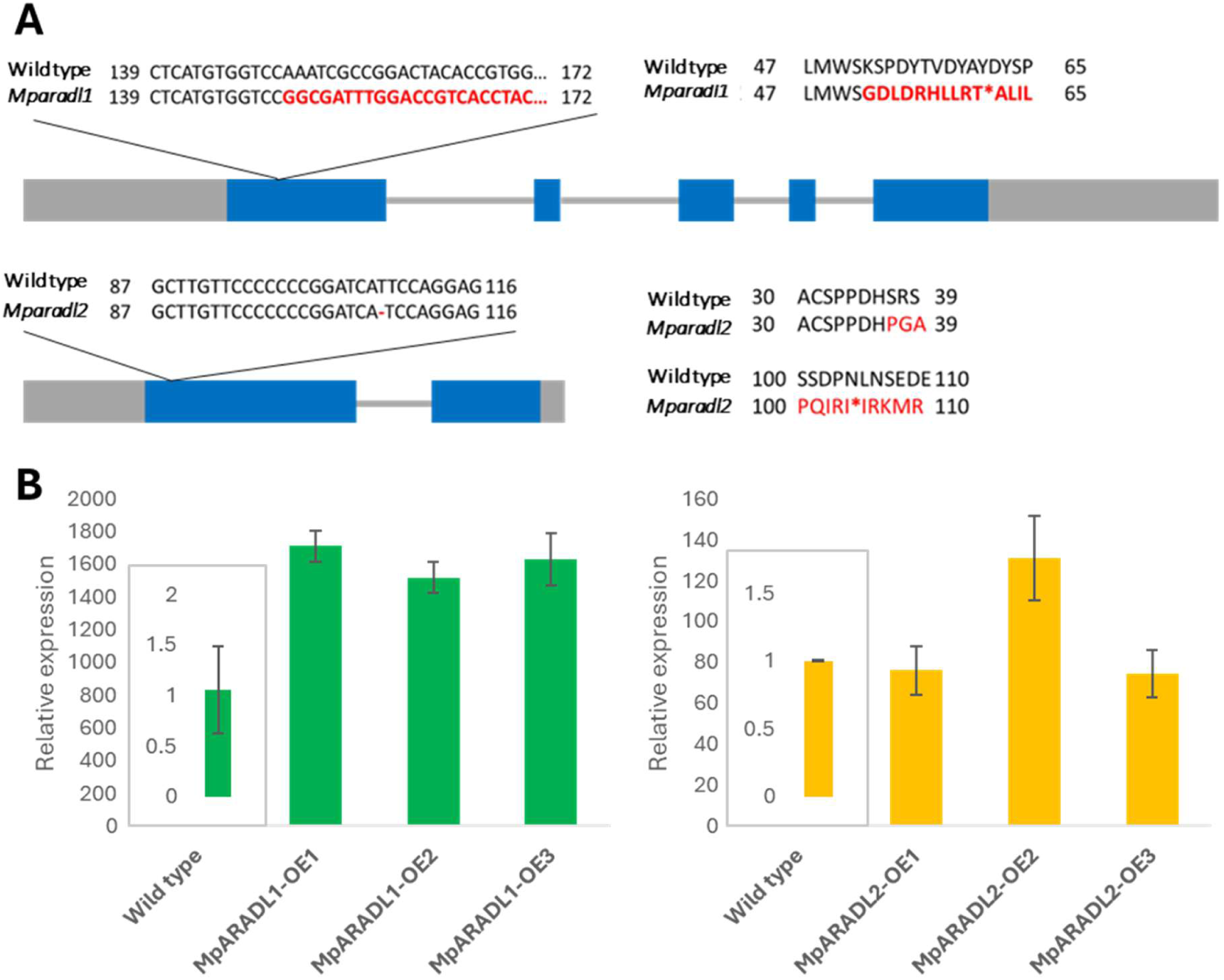
A) Schematic of the gene structures of MpARADL1 (top) and MpARADL2 (bottom). Grey box = untranslated region, blue box = exon, grey lines = introns. Changes in the nucleotide sugar sequence is shown on the left, with red text marking changes observed in the mutant. The resulting changes to the protein sequence is shown on the right. Asterisk shows the stop codon. B) Overexpression of MpARADL1 (green) and MpARADL2 (yellow) was confirmed at a transcript level using qRT-PCR.

**Figure 7.**
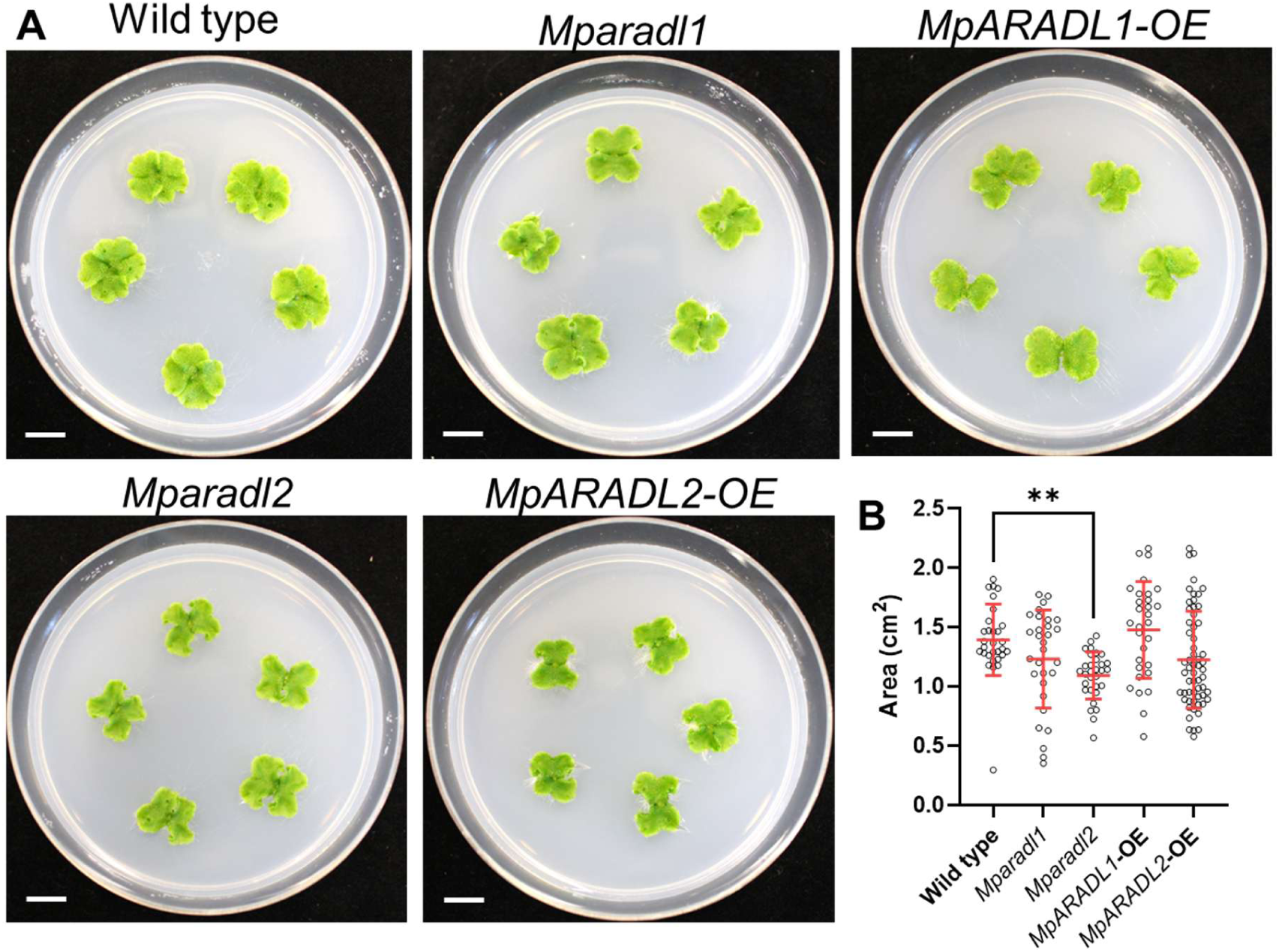
A) Images of 2-week-old plants of MpARADL transgenic lines. Scale bar = 1 cm. B) Quantification of thallus area in 2-week-old transgenic plants compared to wild type. One-way ANOVA was used to identify significant differences between wild type and the transgenic lines. ** = *p* < 0.01.

**Figure 8.**
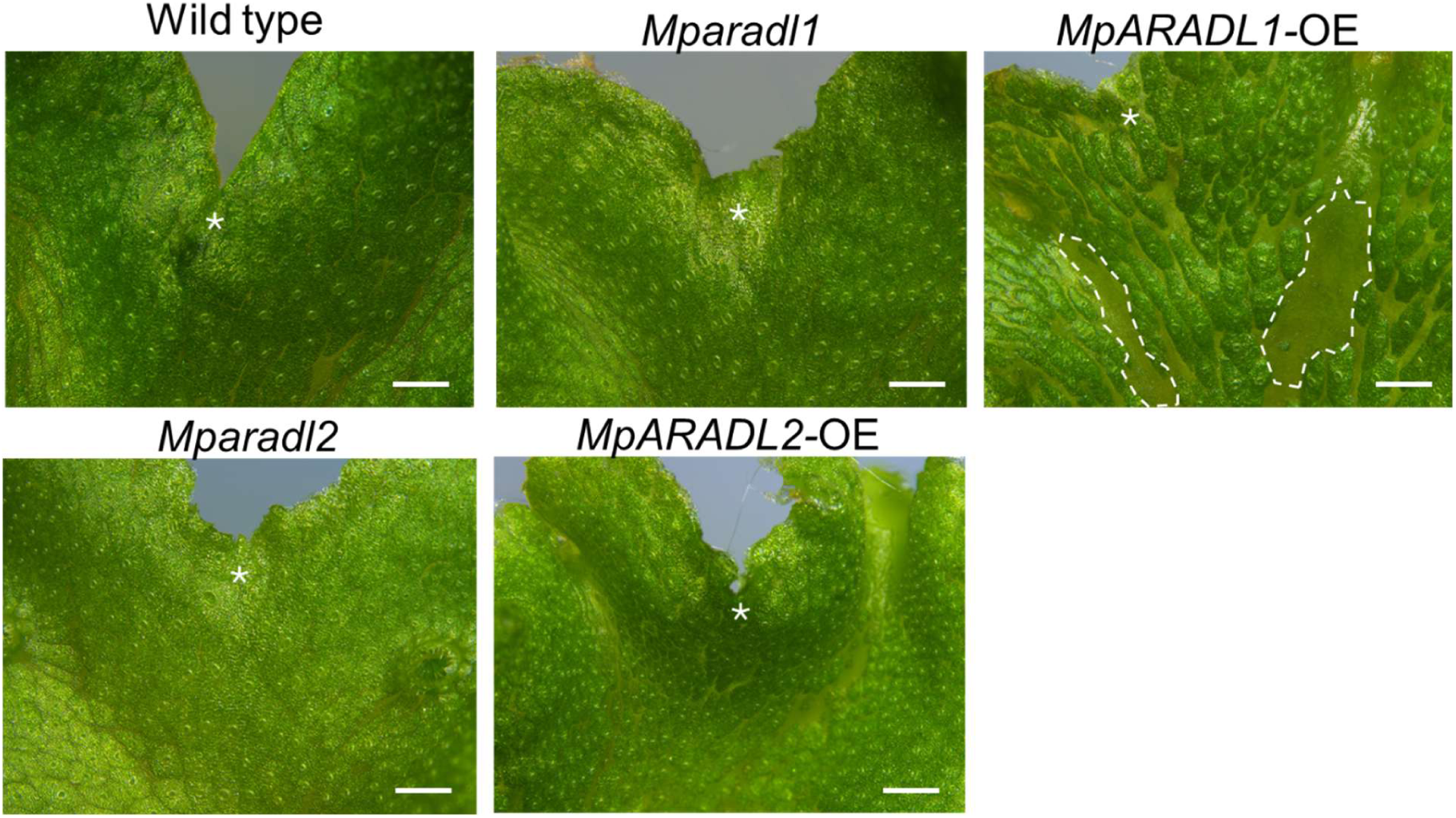
Representative images of the apical notch of 3-week-old Marchantia thalli. All three *MpARADL1* overexpression lines showed a phenotype where the air chambers were not as compactly organised compared to the wild type, often forming empty gaps (dashed lines) on the thallus. Scale bar = 100 µm.

**Figure 9.**
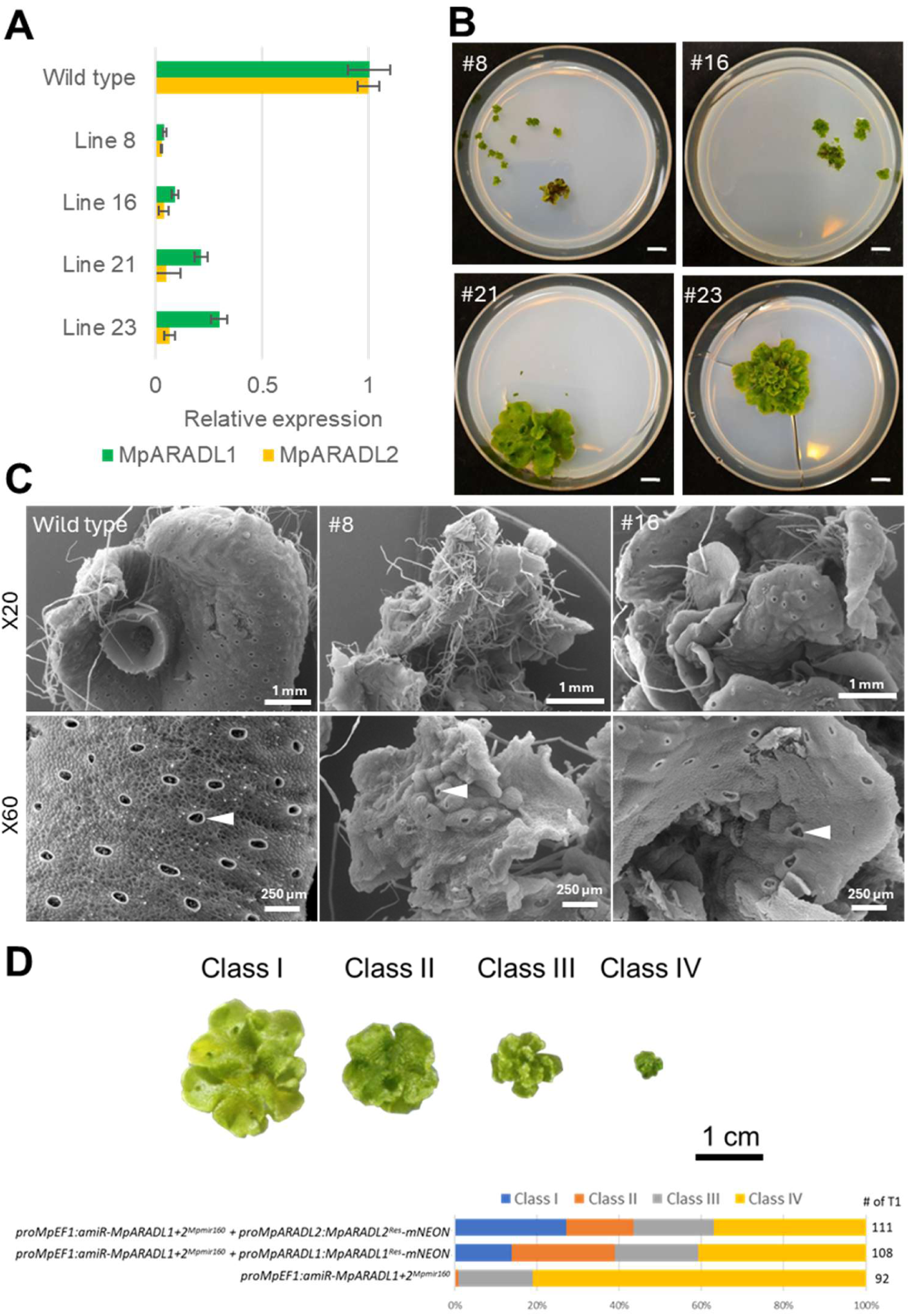
A) Expression of *MpARADL1* and *MpARADL2* in four independent lines expressing *amiR-MpARADL1+2^Mpmir160^* analysed by qRT-PCR. *MpARADL1* and *2* levels are simultaneously downregulated in all mutants, with *MpARADL2* down-regulation being slightly more effective. B) Images of T1 transformants 1 month after transferring to new antibiotic selection media. C) Scanning electron microscope (SEM) images of 2-month old wild type thallus compared to approximately 2-months old #8 and #16 lines show ectopic growth of rhizoids and malformed thalli in the mutants. D) Constructs carrying amiRNA resistant copies of *MpARADL1* or *2* were co-transformed with the amiRNA construct, and the resulting transformant phenotypes were classified into four classes in 3-week-old T1 plants.

### CoMPP and methylation analysis show distinct changes in *MpARADL* transgenic lines

To understand the changes occurring in the cell walls of the MpARADL transgenic plants at a molecular level, we analysed the loss-of-function mutants and OE lines with the Comprehensive Microarray based Polymer Profiling (CoMPP) method (Moller et al. 2007; Pedersen et al. 2012) using 25 cell wall-directed antibodies (Table 1). To investigate changes in specific components of the cell wall, two fractions were sequentially extracted from de-starched AIR using CDTA (trans-1,2-diaminocyclohexane-N,N,N′,N′-tetraacetic acid) and NaOH. CDTA was used to displace cell wall polymers bound through ionic bonds (Jarvis 1982), and NaOH was used to extract polymers more strongly bound to the cell wall by disrupting hydrogen bonds through alkalinisation (Okano and Sarko 1985). Most cell wall-directed antibodies were developed using spermatophyte antigens. Thus, the CoMPP analysis of the *Marchantia* thalli also gives us a useful overview of which cell wall epitopes are also present in *Marchantia* (Table 1). It should be noted that the epitopes recognised by these antibodies may be present in other tissues of Marchantia, such as LM13 in the sporophytes (Kang et al., 2025). It is also possible that the signal was below the threshold for detection and therefore does not necessarily preclude the presence of these cell wall epitopes in the Marchantia thallus.

**Table 1.** Antibodies used for Comprehensive Microarray based Polymer Profiling (CoMPP). 0 = not detected, 1-10 (+), 11-30 (++), 30+ (+++).

| Antibody | Epitope | Reference | Detection in wild type<br>Marchantia thalli |  |
| --- | --- | --- | --- | --- |
|  |  |  | CDTA | NaOH |
| Pectin |  |  |  |  |
| LM5 | β-(1,5)-galactan | (Andersen et al., 2016; Ruprecht et al., 2017) | n.d. | n.d. |
| LM6 | α-(1,5)-arabinan | (Verhertbruggen, Marcus, Haeger, Verhoef, et al., 2009) | n.d. | + |
| LM13 | α-(1,5)-arabinan | (Verhertbruggen, Marcus, Haeger, Verhoef, et al., 2009) | n.d. | n.d. |
| LM16 | Galactosylated RG-I | (Pedersen et al. 2012; Verhertbruggen, et al.,2009) | ++ | ++ |
| LM18 | Partially methyl esterified homogalacturonan | (Verhertbruggen, Marcus, Haeger, Ordaz-Ortiz, et al., 2009) | +++ | + |
| LM19 | Unesterified homogalacturonan | (Verhertbruggen, Marcus, Haeger, Ordaz-Ortiz, et al., 2009) | +++ | + |
| LM20 | Methyl esterified homogalacturonan | (Verhertbruggen, Marcus, Haeger, Ordaz-Ortiz, et al., 2009) | +++ | n.d. |
| INRA-RU1 | >6 Rha-(1,4)-GalA-(1,2) disaccharide backbone repeats of RG-I | (Ralet et al., 2010) | + | ++ |
| INRA-RU2 | >2-7 Rha-(1,4)-GalA-(1,2) disaccharide backbone repeats of RG-I | (Ralet et al., 2010) | + | ++ |
| Glycoprotein |  |  |  |  |
| JIM8 | Arabinogalactan | (Pennell et al., 1991) | + | ++ |
| JIM13 | (1,3) -β-GlcA-(1,3)-α-GalA-(1,2)-Rha in AGPs | (Pennell et al., 1991) | +++ | +++ |
| JIM15 | β-GlcA | (Yates et al., 1996) | n.d. | n.d. |
| LM1 | Extensins | (Smallwood, Martin, and Knox 1995) | n.d. | ++ |
| LM2 | β-GlcA | (Smallwood et al. 1995) | ++ | +++ |
| Hemicellulose |  |  |  |  |
| **LM8** | Xylogalacturonan | (Willats et al., 2004) | n.d. | n.d. |
| **LM10** | β-(1,4)-Xyl | (Pedersen et al. 2012) | n.d. | n.d. |
| **LM11** | Arabinoxylan | (Pedersen et al. 2012) | n.d. | + |
| **LM15** | Xyloglucan XXXG motif | (Marcus et al. 2008; Pedersen et al. 2012) | + | + |
| **LM21** | (1,4)-β-mannan | (Marcus et al. 2010; Pedersen et al. 2012) | ++ | +++ |
| **LM23** | (1,3)-β-xylose residues in xylan | (Manabe et al. 2011) | + | + |
| **LM24** | Xyloglucan XLLG motif | (Pedersen et al. 2012) | n.d. | n.d. |
| **LM25** | Xyloglucan XXXG/XLXG/XLLG motifs and linear (1,4)-β-linked glucopentose | (Pedersen et al. 2012) | +++ | +++ |
| **CCRC-M1** | Fucosylated xyloglucan α-Fuc-β-Gal | (Pattathil et al. 2010) | n.d. | n.d. |
| **CCRC-M39** | Fucosylated xyloglucan and RG-I from sycamore plants | (Pattathil et al. 2010) | n.d. | + |
| **CCRC-M58** | Xyloglucan XLLG motif | (Pattathil et al. 2010) | n.d. | n.d. |

The antibody binding for *Mparadl1* was mostly comparable to wild type, except for reduced labelling of the LM2 (β-GlcA) AGP antibody (Figure 10). *MpARADL1* OE lines consistently showed increased binding of the LM25 (xyloglucan) antibody. *MpARADL1* OE lines also showed reduced binding of RG-I backbone directed antibodies, INRA-RU1 (Figure 10). The *Mparadl2* antibody binding patterns showed reduced LM16 (galactosylated RG-I) epitope, and increased LM2 and INRA-RU1 epitopes. In the *MpARADL2* OE line, INRA-RU1 labelling was abolished, contrary to the increased detection of these antibodies in *Mparadl2*. The distinctive changes in the cell wall epitope binding support the non-redundant roles for *MpARADL1* and *2*, but both genes appear to affect RG-I or polymers associated with it. The amiRNA mutants could unfortunately not be tested using CoMPP due to its aberrant phenotype, and the requirement of 10 mg of Alcohol Insoluble Residue (AIR) material per technical replicate for analysis.

**Figure 10.**
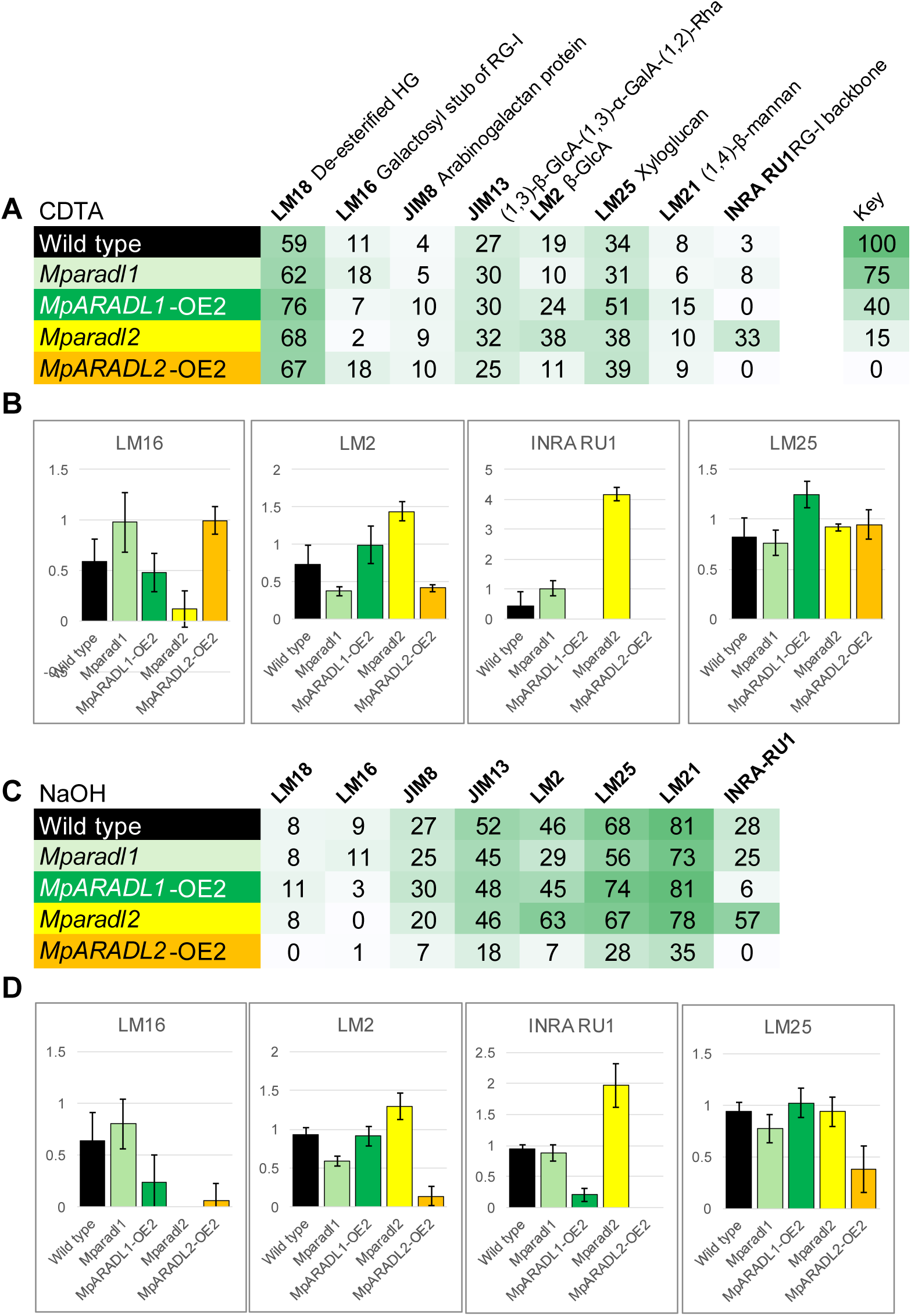
Comprehensive Microarray based Polymer Profiling (CoMPP) results for *MpARADL* transgenic lines displayed in a heatmap for select antibodies. Results are an average of six technical replicates. Heatmap for the CDTA (A) and NaOH (D) fractions. The highest values recorded in wild type for the CDTA (B) or NaOH fractions (D) were used to calculate relative detection levels across all replicates for four antibodies. C) Heatmap for the NaOH fraction.

To investigate whether the results from CoMPP could be explained by changes in the glycosyl linkage composition of the *MpARADL* mutants, the transgenic lines were analysed using methylation analysis (Table 2). The glycosyl linkages associated with arabinan, and with epitopes of the antibodies that showed differential binding in the transgenic lines, including LM16, LM2, INRA-RU1 and LM25 are shown in Figure 11. The 5-linked Ara*f* linkages, which is associated with arabinan, were comparable across all transgenic lines and wild type samples at low levels (∼-0.5 mol%) (Figure 11A). This indicates that there are likely other GTs responsible for the 5-linked Ara*f* linkages in the Marchantia thallus. The low relative 5-Ara*f* content is consistent with the CoMPP data which showed very low detection of the LM6 epitope in the NaOH fraction of wild type plants. The Terminal-Ara*f* (t-Ara*f*) linkage, which is also associated with arabinan, was slightly elevated in *Mparadl1* and the amiRNA mutants. Overall, the modulation *MpARADL1* and *2* expressions did not lead to changes in arabinan-related glycosyl linkages.

**Table 2.**
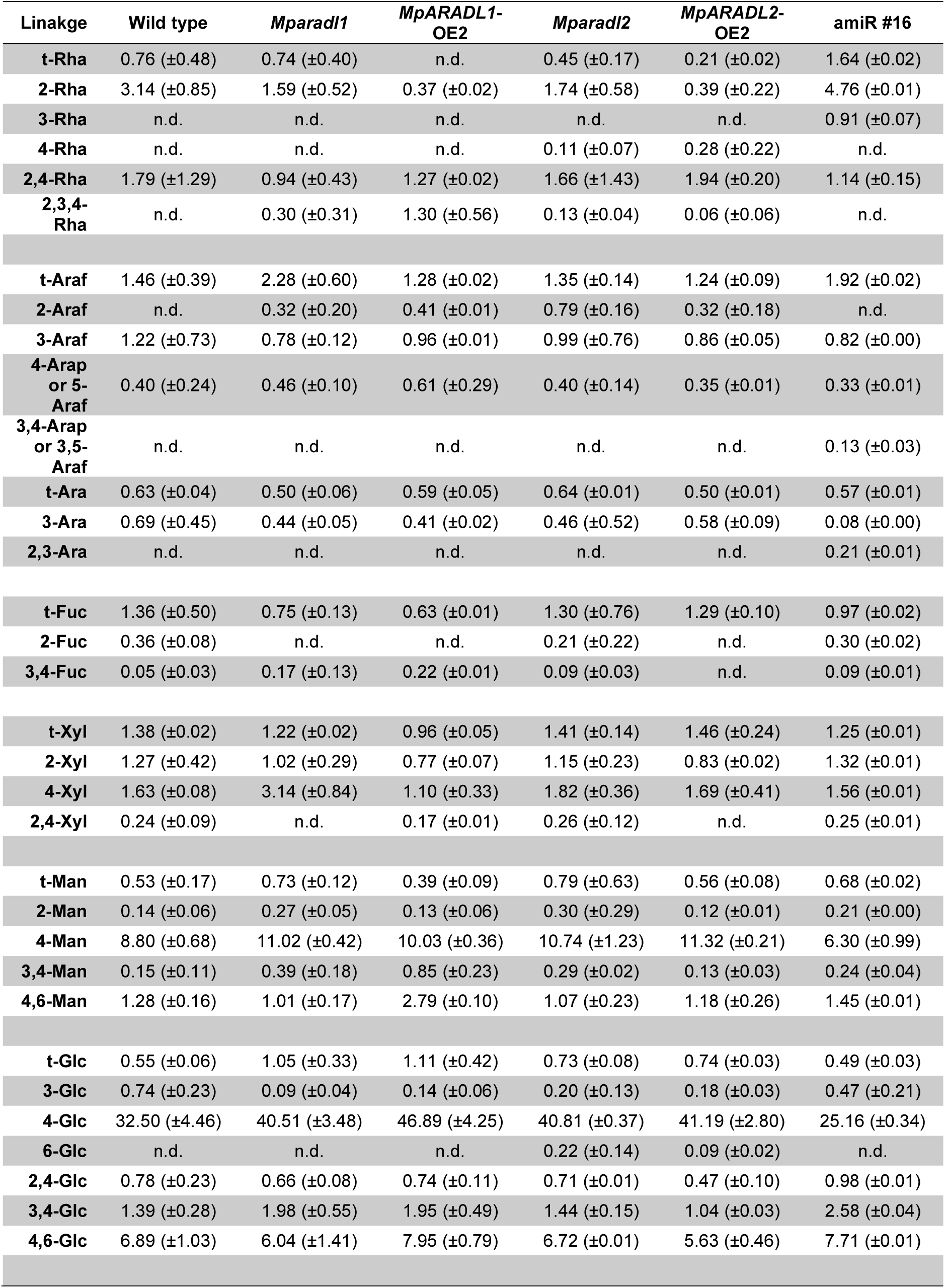
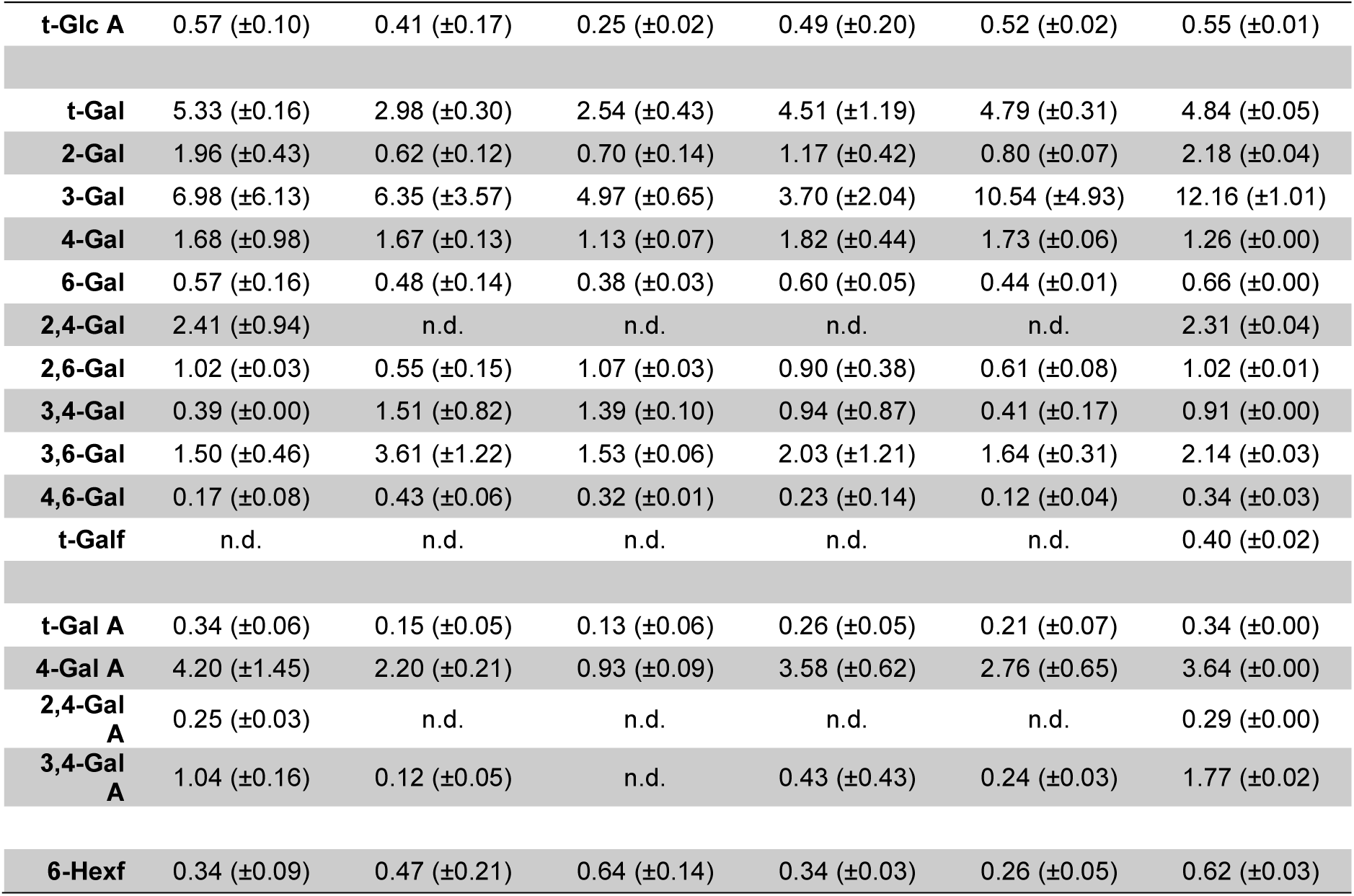
Glycosyl linkage analysis of *MpARADL* transgenic lines. n=2. n.d. = not detected. Standard deviation values are shown in brackets.

| Linakge | Wild type | *Mparadl1* | *MpARADL1*-<br>OE2 | *Mparadl2* | *MpARADL2*-<br>OE2 | amiR #16 |
| --- | --- | --- | --- | --- | --- | --- |
| **t-Rha** | 0.76 (±0.48) | 0.74 (±0.40) | n.d. | 0.45 (±0.17) | 0.21 (±0.02) | 1.64 (±0.02) |
| **2-Rha** | 3.14 (±0.85) | 1.59 (±0.52) | 0.37 (±0.02) | 1.74 (±0.58) | 0.39 (±0.22) | 4.76 (±0.01) |
| **3-Rha** | n.d. | n.d. | n.d. | n.d. | n.d. | 0.91 (±0.07) |
| **4-Rha** | n.d. | n.d. | n.d. | 0.11 (±0.07) | 0.28 (±0.22) | n.d. |
| **2,4-Rha** | 1.79 (±1.29) | 0.94 (±0.43) | 1.27 (±0.02) | 1.66 (±1.43) | 1.94 (±0.20) | 1.14 (±0.15) |
| **2,3,4-Rha** | n.d. | 0.30 (±0.31) | 1.30 (±0.56) | 0.13 (±0.04) | 0.06 (±0.06) | n.d. |
| **t-Araf** | 1.46 (±0.39) | 2.28 (±0.60) | 1.28 (±0.02) | 1.35 (±0.14) | 1.24 (±0.09) | 1.92 (±0.02) |
| **2-Araf** | n.d. | 0.32 (±0.20) | 0.41 (±0.01) | 0.79 (±0.16) | 0.32 (±0.18) | n.d. |
| **3-Araf** | 1.22 (±0.73) | 0.78 (±0.12) | 0.96 (±0.01) | 0.99 (±0.76) | 0.86 (±0.05) | 0.82 (±0.00) |
| **4-Araf**<br>**or 5-Araf** | 0.40 (±0.24) | 0.46 (±0.10) | 0.61 (±0.29) | 0.40 (±0.14) | 0.35 (±0.01) | 0.33 (±0.01) |
| **3,4-Araf**<br>**or 3,5-Araf** | n.d. | n.d. | n.d. | n.d. | n.d. | 0.13 (±0.03) |
| **t-Ara** | 0.63 (±0.04) | 0.50 (±0.06) | 0.59 (±0.05) | 0.64 (±0.01) | 0.50 (±0.01) | 0.57 (±0.01) |
| **3-Ara** | 0.69 (±0.45) | 0.44 (±0.05) | 0.41 (±0.02) | 0.46 (±0.52) | 0.58 (±0.09) | 0.08 (±0.00) |
| **2,3-Ara** | n.d. | n.d. | n.d. | n.d. | n.d. | 0.21 (±0.01) |
| **t-Fuc** | 1.36 (±0.50) | 0.75 (±0.13) | 0.63 (±0.01) | 1.30 (±0.76) | 1.29 (±0.10) | 0.97 (±0.02) |
| **2-Fuc** | 0.36 (±0.08) | n.d. | n.d. | 0.21 (±0.22) | n.d. | 0.30 (±0.02) |
| **3,4-Fuc** | 0.05 (±0.03) | 0.17 (±0.13) | 0.22 (±0.01) | 0.09 (±0.03) | n.d. | 0.09 (±0.01) |
| **t-Xyl** | 1.38 (±0.02) | 1.22 (±0.02) | 0.96 (±0.05) | 1.41 (±0.14) | 1.46 (±0.24) | 1.25 (±0.01) |
| **2-Xyl** | 1.27 (±0.42) | 1.02 (±0.29) | 0.77 (±0.07) | 1.15 (±0.23) | 0.83 (±0.02) | 1.32 (±0.01) |
| **4-Xyl** | 1.63 (±0.08) | 3.14 (±0.84) | 1.10 (±0.33) | 1.82 (±0.36) | 1.69 (±0.41) | 1.56 (±0.01) |
| **2,4-Xyl** | 0.24 (±0.09) | n.d. | 0.17 (±0.01) | 0.26 (±0.12) | n.d. | 0.25 (±0.01) |
| **t-Man** | 0.53 (±0.17) | 0.73 (±0.12) | 0.39 (±0.09) | 0.79 (±0.63) | 0.56 (±0.08) | 0.68 (±0.02) |
| **2-Man** | 0.14 (±0.06) | 0.27 (±0.05) | 0.13 (±0.06) | 0.30 (±0.29) | 0.12 (±0.01) | 0.21 (±0.00) |
| **4-Man** | 8.80 (±0.68) | 11.02 (±0.42) | 10.03 (±0.36) | 10.74 (±1.23) | 11.32 (±0.21) | 6.30 (±0.99) |
| **3,4-Man** | 0.15 (±0.11) | 0.39 (±0.18) | 0.85 (±0.23) | 0.29 (±0.02) | 0.13 (±0.03) | 0.24 (±0.04) |
| **4,6-Man** | 1.28 (±0.16) | 1.01 (±0.17) | 2.79 (±0.10) | 1.07 (±0.23) | 1.18 (±0.26) | 1.45 (±0.01) |
| **t-Glc** | 0.55 (±0.06) | 1.05 (±0.33) | 1.11 (±0.42) | 0.73 (±0.08) | 0.74 (±0.03) | 0.49 (±0.03) |
| **3-Glc** | 0.74 (±0.23) | 0.09 (±0.04) | 0.14 (±0.06) | 0.20 (±0.13) | 0.18 (±0.03) | 0.47 (±0.21) |
| **4-Glc** | 32.50 (±4.46) | 40.51 (±3.48) | 46.89 (±4.25) | 40.81 (±0.37) | 41.19 (±2.80) | 25.16 (±0.34) |
| **6-Glc** | n.d. | n.d. | n.d. | 0.22 (±0.14) | 0.09 (±0.02) | n.d. |
| **2,4-Glc** | 0.78 (±0.23) | 0.66 (±0.08) | 0.74 (±0.11) | 0.71 (±0.01) | 0.47 (±0.10) | 0.98 (±0.01) |
| **3,4-Glc** | 1.39 (±0.28) | 1.98 (±0.55) | 1.95 (±0.49) | 1.44 (±0.15) | 1.04 (±0.03) | 2.58 (±0.04) |
| **4,6-Glc** | 6.89 (±1.03) | 6.04 (±1.41) | 7.95 (±0.79) | 6.72 (±0.01) | 5.63 (±0.46) | 7.71 (±0.01) |
| **t-Glc A** | 0.57 (±0.10) | 0.41 (±0.17) | 0.25 (±0.02) | 0.49 (±0.20) | 0.52 (±0.02) | 0.55 (±0.01) |
| **t-Gal** | 5.33 (±0.16) | 2.98 (±0.30) | 2.54 (±0.43) | 4.51 (±1.19) | 4.79 (±0.31) | 4.84 (±0.05) |
| **2-Gal** | 1.96 (±0.43) | 0.62 (±0.12) | 0.70 (±0.14) | 1.17 (±0.42) | 0.80 (±0.07) | 2.18 (±0.04) |
| **3-Gal** | 6.98 (±6.13) | 6.35 (±3.57) | 4.97 (±0.65) | 3.70 (±2.04) | 10.54 (±4.93) | 12.16 (±1.01) |
| **4-Gal** | 1.68 (±0.98) | 1.67 (±0.13) | 1.13 (±0.07) | 1.82 (±0.44) | 1.73 (±0.06) | 1.26 (±0.00) |
| **6-Gal** | 0.57 (±0.16) | 0.48 (±0.14) | 0.38 (±0.03) | 0.60 (±0.05) | 0.44 (±0.01) | 0.66 (±0.00) |
| **2,4-Gal** | 2.41 (±0.94) | n.d. | n.d. | n.d. | n.d. | 2.31 (±0.04) |
| **2,6-Gal** | 1.02 (±0.03) | 0.55 (±0.15) | 1.07 (±0.03) | 0.90 (±0.38) | 0.61 (±0.08) | 1.02 (±0.01) |
| **3,4-Gal** | 0.39 (±0.00) | 1.51 (±0.82) | 1.39 (±0.10) | 0.94 (±0.87) | 0.41 (±0.17) | 0.91 (±0.00) |
| **3,6-Gal** | 1.50 (±0.46) | 3.61 (±1.22) | 1.53 (±0.06) | 2.03 (±1.21) | 1.64 (±0.31) | 2.14 (±0.03) |
| **4,6-Gal** | 0.17 (±0.08) | 0.43 (±0.06) | 0.32 (±0.01) | 0.23 (±0.14) | 0.12 (±0.04) | 0.34 (±0.03) |
| **t-Galf** | n.d. | n.d. | n.d. | n.d. | n.d. | 0.40 (±0.02) |
| **t-Gal A** | 0.34 (±0.06) | 0.15 (±0.05) | 0.13 (±0.06) | 0.26 (±0.05) | 0.21 (±0.07) | 0.34 (±0.00) |
| **4-Gal A** | 4.20 (±1.45) | 2.20 (±0.21) | 0.93 (±0.09) | 3.58 (±0.62) | 2.76 (±0.65) | 3.64 (±0.00) |
| **2,4-Gal A** | 0.25 (±0.03) | n.d. | n.d. | n.d. | n.d. | 0.29 (±0.00) |
| **3,4-Gal A** | 1.04 (±0.16) | 0.12 (±0.05) | n.d. | 0.43 (±0.43) | 0.24 (±0.03) | 1.77 (±0.02) |
| **6-Hexf** | 0.34 (±0.09) | 0.47 (±0.21) | 0.64 (±0.14) | 0.34 (±0.03) | 0.26 (±0.05) | 0.62 (±0.03) |

**Figure 11.**
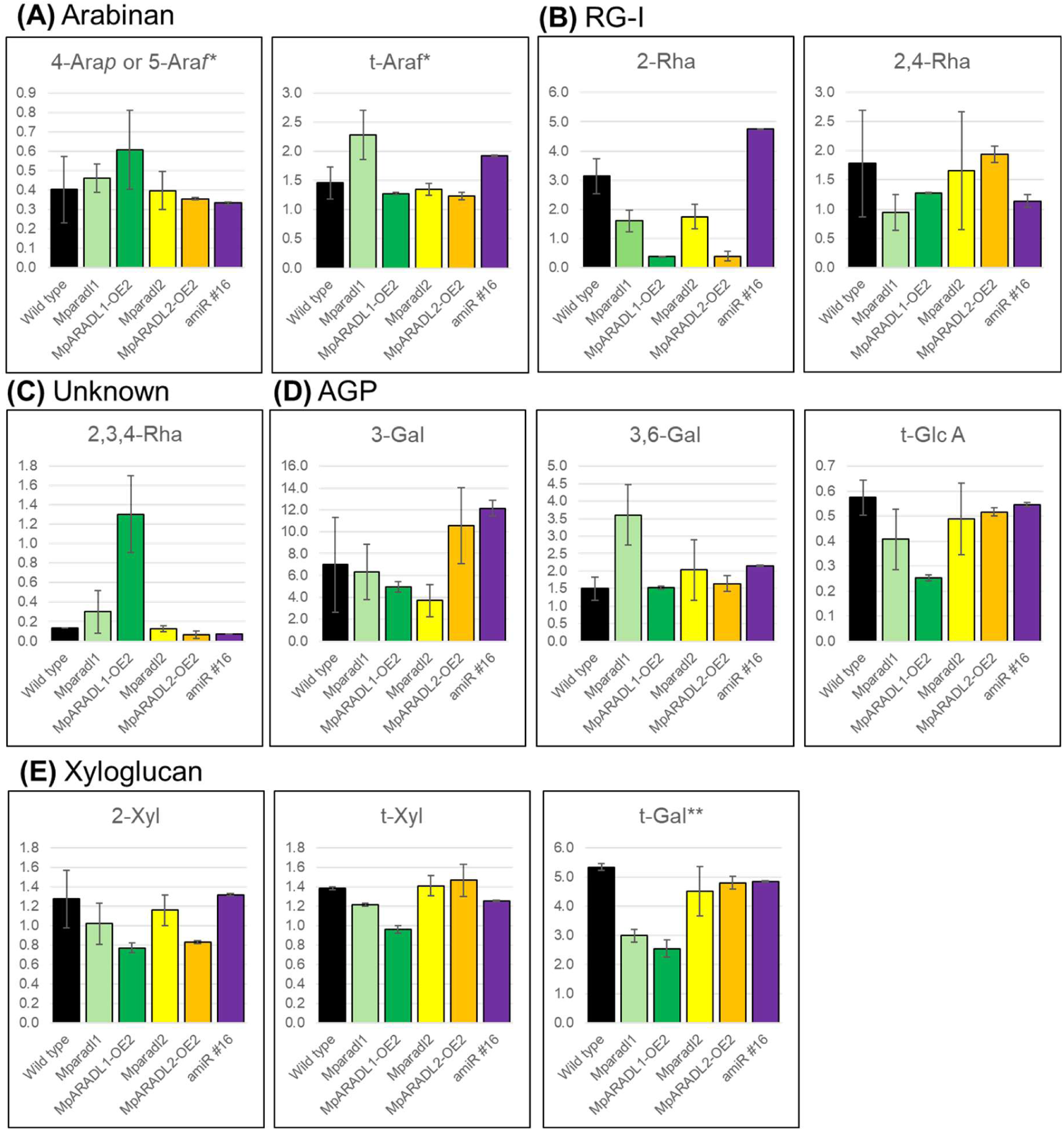
Select glycosyl linkages from the methylation analysis related to (A) arabinan, and epitope of (B) INRA-RU1 (RG-I backbone), (C) an unknown polymer, (D) epitope of LM2 (AGP β-GlcA) and AGPs, and (E) epitopes of LM25 (Xyloglucan XXXG/XLXG/XLLG motifs). * = also related to RG-I and AGPs. **= Also related to LM16 (Galactosylated RG-I) epitopes. Values are represented in mol%.

2-Rha (Rhamnose) and 2,4-Rha are linkages associated with the backbone of RG-I. RG-I has a backbone of [4-α-GalA-(1,2)-α-Rhap-1] disaccharide repeats (Renard et al. 1997) synthesised by RG-I RHAMNOSYLTRANSFERASES (RRT) in GT family 106 and RG-I: Galaturonosyltransferase 1 (RGGAT1) in GT family 116 (Amos et al. 2022; Wachananawat et al. 2020). The rhamnosyl residue of the RG-I backbone can be branched at the *O-*4 position with linear or branched versions of (1,5)-α-arabinan and (1,4)-β-galactose polymers (galactan). Interestingly, 2-Rha levels were reduced in *Mparadl1*, *Mparadl2*, *MpARADL1*-OE and *MpARADL2*-OE, suggesting that the activity of MpARADL1 and 2 may influence RG-I backbone biosynthesis (Figure 11B). Interestingly, 2,3,4-Rha linkages were also detected in the methylation analysis. 2,3,4-linked Rha residue is associated with RG-II in *Arabidopsis,* which has an apiose monosaccharide attached to the first anomeric carbon of the 2,3,4-linked Rha (Bar-Peled, Urbanowicz, and O’Neill 2012). To date, there is no definitive evidence for the presence of RG-II in bryophytes (Pfeifer et al., 2022). Furthermore, overexpression of the UDP-Apiose synthase (UAS) which biosynthesises UDP-apiose, a substrate of apiosyltransferases did not lead to apiose incorporated into the *P. patens* cell wall (Smith et al. 2016). Thus, the 2,3,4-Rha linkage in Marchantia is likely not associated with RG-II, but with another polymer. In the *MpARADL1* OE line, there was almost a 10-fold increase in the relative amount of 2,3,4-Rha (Figure 11C). In regard to the antibody binding pattern of INRA-RU1, perhaps 2,3,4-Rha could be a part of RG-I in Marchantia and the increased branching of the Rha residues in *MpARADL1*-OE could be hindering the antibody access to the RG-I backbone. The 2,3,4-Rha residue is still present in the *Mparadl1* mutant, which is consistent with the CoMPP results of *Mparadl1* lines showing similar INRA-RU epitope recognition as in wild type. Hence, there could be an indirect influence of *MpARADL1* activity on 2,3,4-Rha content rather than a direct link to its catalytic function.

Next, we analysed the linkages associated with type II arabinogalactan (type II AG) sidechains of AGPs which harbour terminal glucuronic acid (t-GlcA) residues that bind LM2 antibody (Figure 11D). Type II AG is a (1,3)-β-galactan branched with galactosyl residues at the *O*-6 position, which can be further elongated into a (1,6)-β-galactan chain. The *O*-6 galactosyl moieties of type II AG can be terminated with other sugars, such as arabinose, methylated glucuronic acid and fucose (Haque et al. 2005; Tryfona et al. 2014; Soto et al. 2021). In *Arabidopsis*, t-GlcA of type II AG can be covalently linked with the Rha residues of the RG-I backbone (Tan et al. 2022). The 3-Gal (galactose) linkage, which is catalysed by KAONASHI (KNS4) in GT family 31 (Suzuki et al. 2017), was elevated in *MpARADL2*-OE and reduced in the *Mparadl2* mutant, which is inversely correlated to the binding pattern of LM2. The t-GlcA linkage content, which is the epitope detected by LM2, was comparable between wild type, *Mparadl2* and *MpARADL2*-OE. Therefore, the changes in LM2 labelling do not appear to be a direct result of changes in epitope abundance, but rather its accessibility by antibodies within the cell wall. CoMPP results showed elevated LM25 antibody binding in *MpARADL1*-OE. However, the xyloglucan LM25 epitope related glycosyl linkages, 2-Xyl (xylose), t-Xyl and t-Gal were not elevated in *MpARADL1*-OE (Figure 11E), also indicating that changed antibody binding is likely an artifact of altered antibody accessibility in the cell wall, rather than changes in the percentage of linkages associated with the epitope.

### Immunolabelling reveals potential role of *Mparadl2* in arabinan biosynthesis in elaters

As methylation analysis showed that the thallus contains low amounts of arabinan, we probed the changes in the level of arabinan in the sporophytes, which contain high amounts of arabinan associated with elaters (Kang et al., 2025, Dierschke et al. 2024). We conducted immunolabelling on the resin-embedded sections of mature sporophytes of *Mparadl1* and *Mparadl2* (Figure 12). Calcofluor white, which stains cellulose was used to observe the morphology of the elaters. The LM25 xyloglucan antibody was used as a positive control, as LM25 binding is abundant in the sporophytic cell walls of transfer cells (Henry, Lopez, and Renzaglia 2020). LM25 showed consistent binding when elaters were probed across all mutants. Strikingly, *Mparadl2* showed diminished binding when probed with the LM6 arabinan antibody. As the sporophyte cells are diploid, the residual arabinan labelling could be accounted for by the wild-type male copy of *MpARADL2*. Elaters are specialised cells found in sporophytes, the diploid phase of the *Marchantia* life cycle. They differentiate from archesporial cells into elaterocytes, which undergoe secondary cell wall deposition and programmed cell death (Kremer and Drinnan 2003; Shimamura 2016). During sporophyte maturation, the spore capsule undergoes dehiscence, and elaters take a helical shape upon drying. When the spore capsule bursts, elaters exhibit a rapid twisting motion that aids in spore dispersal. As the structure of *Mparadl2* elaters is comparable to wild type, it appears arabinan may not be essential during the development of elaters. However, it is tempting to speculate that the loss of the 1,5-α-L-arabinan epitope in *Mparadl2* elaters leads to physiological consequences such as reduced hygroscopic movements. In sugar beet and potato pectin, arabinan side chains show high levels of flexibility and interacts readily with water molecules (Ha et al. 2005; Larsen et al. 2011). Whether arabinan is important for harnessing moisture that ultimately facilitates the hygroscopic movement of elaters requires further investigation.

**Figure 12.**
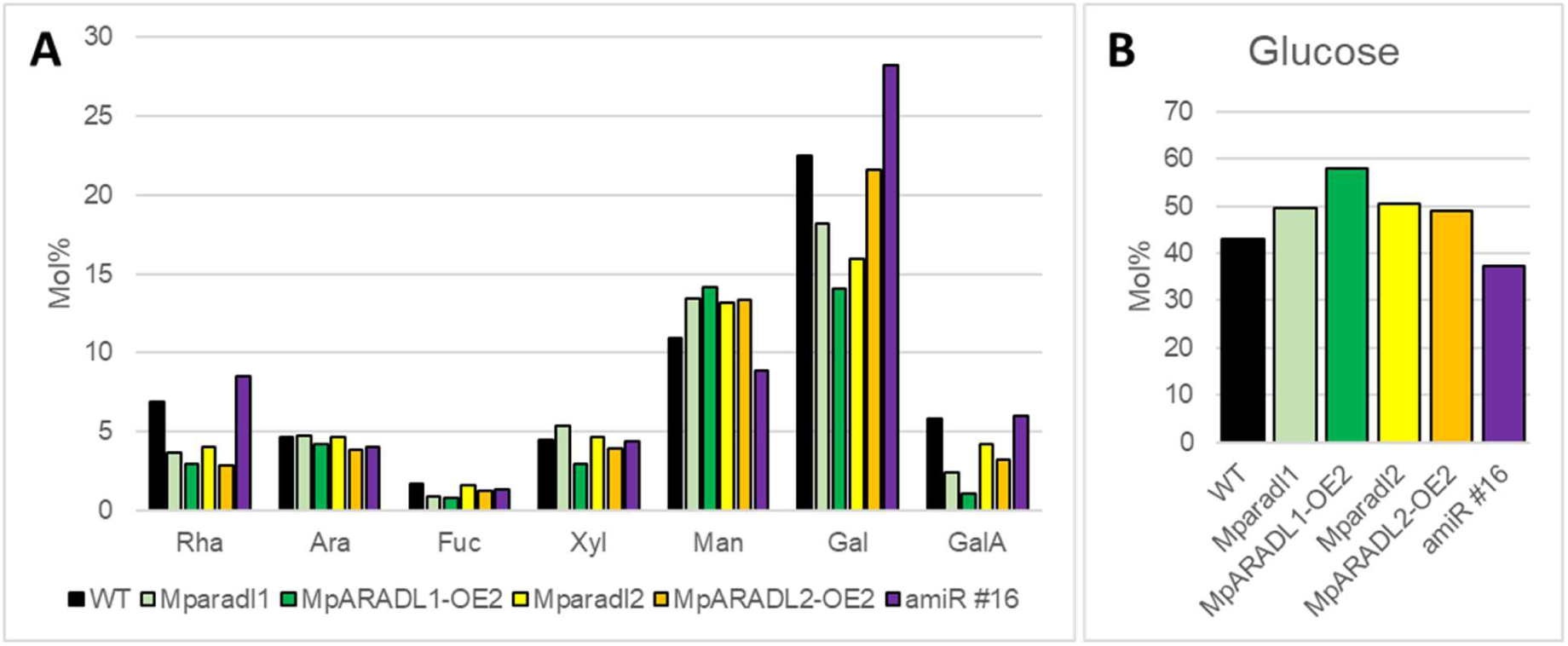
A) Total relative mol% of eight different monosaccharides deduced from the glycosyl linkage analysis. Values for glucose (B) are shown separately.

**Figure 12.**
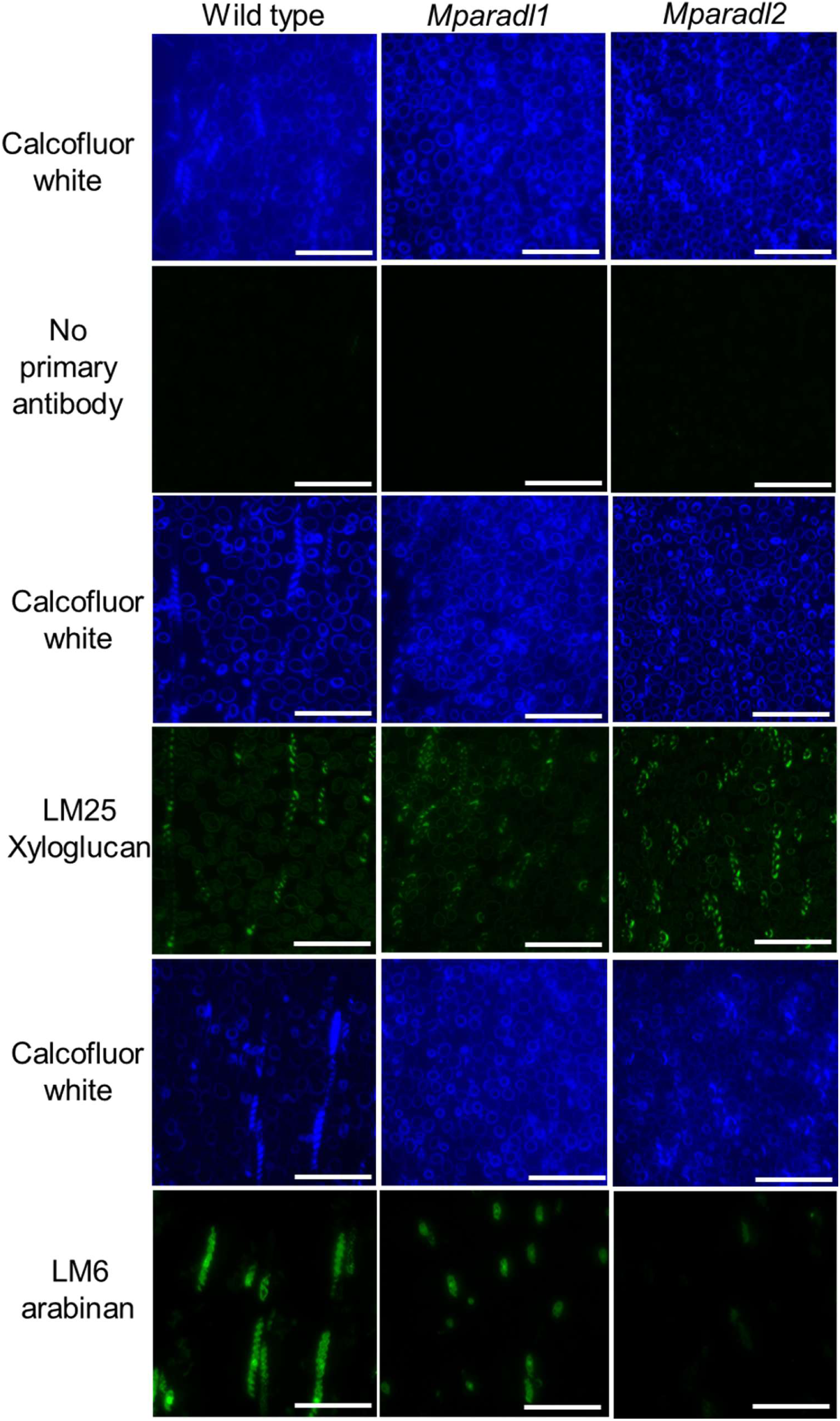
Images from resin embedded sections of mature spore capsule of Marchantia labelled with calcofluor white, no primary antibody, LM25 xyloglucan antibody and LM6 α-1,5-arabinan antibody. Scale bar 50 µm.

### GT47 clade B proteins are difficult to purify in the HEK293 cell heterologous expression system

The establishment of the HEK293 cell heterologous expression system for cell wall biosynthetic GTs has led to the successful expression and purification of multiple plant glycoenzymes for downstream catalytic activity assays (Amos et al. 2018; Prabhakar et al. 2020; Ruprecht et al. 2020; Soto et al. 2021). In this study, we expressed the catalytic domains of AtARAD1, as well as MpARADL1 and MpARADL2 in the HEK293 cell heterologous expression system. The transmembrane region of proteins was predicted using TOPCONS (https://topcons.cbr.su.se/pred/; Tsirigos et al. 2015) (Figure 13A), and the N-terminus of the coding sequence was truncated at the beginning of the predicted luminal region. Further troubleshooting efforts to avoid potential N-glycosylation sites at the beginning of the luminal region led to multiple N-terminally truncated versions of these proteins, but did not lead to soluble proteins (Figure 13B). As AtARAD1 has been shown to interact with other GT47 clade B proteins such as AtARAD2 and the *N. alata* ARADL1, we attempted to co-express AtARAD1 with the Marchantia ARADLs, but the proteins could not be solubilised (Figure 13C).

**Figure 13.**
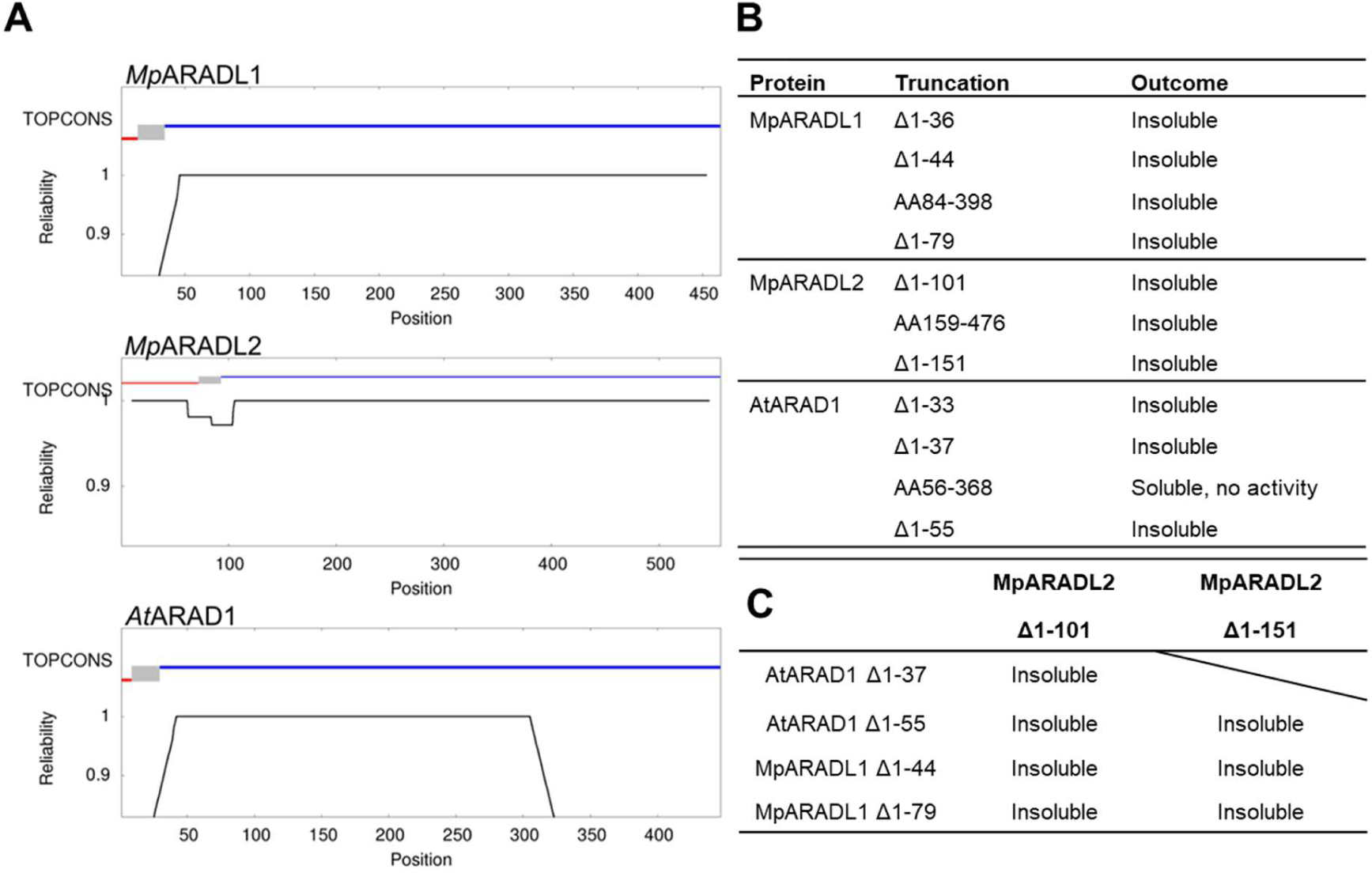
A) TOPCONS predictions of protein topology. Red line indicates the cytosolic N-terminal tail, the grey box indicates the transmembrane region, and the blue line indicates the predicted catalytic domain residing in the Golgi lumen. Amino acid positions are shown on the bottom. B) Description of the truncated catalytic domains expressed in the HEK293 cell system. (C) Description of the truncated catalytic domains co-expressed in the HEK293 cell system. AA = amino acids.

Despite the broad biological roles of arabinans in plant physiology, little is known about the mechanisms underlying their biosynthesis. The biochemical function of putative arabinan GTs has not yet been tested due to difficulties in purifying the enzyme. Previously, AtARAD1 activity was surveyed by using microsomal fractions of *N. benthamiana* plants transiently overexpressing AtARAD1 (Dr. Jesper Harholt pers. comm.). In this study, we attempted to use the HEK293 cell expression system to characterise the precise catalytic activity of putative arabinan GTs *in vitro*. Here we attempted to optimise GT47 clade B proteins by truncating them further along the N-terminus to avoid potential N-glycosylation sites, and co-expressed them with other proteins to assess interaction-dependent conformation changes which can aid with the solubility of GTs (Amos et al. 2018). However, the GT47 clade B proteins tested in this study could not be secreted into the media for subsequent purification steps. Considering that GT47 clade B proteins from two separate embryophyte lineages both exhibit difficulties in purification, it appears that the challenges associated with purifying GT47 clade B proteins are likely not limited by the heterologous expression system or the model organism, but by yet unknown factors which hinders proper protein folding. With the rapid development of in silico modelling of multimeric protein complexes and new tools to assess protein-protein interactions, such as proximity labelling assays, an exploration into a larger catalytic module for arabinan biosynthesis may be necessary for proof of catalytic activity.

### Conclusion

*MpARADL1* and *MpARADL2* are characterised in the liverwort model organism, *M. polymorpha*, with their misexpression leading to distinct changes in cell wall composition. The role of *MpARADL2* in the biosynthesis of arabinan in the elaters is highlighted. The relatively new HEK293 cell heterologous expression system could not be used to isolate soluble AtARAD1 and MpARADL1 and 2 proteins for catalytic activity assays. Our attempts to optimise the constructs using different N-terminal truncations and co-expression of proteins suggest that GT47 clade B proteins require a yet unknown mechanism to confer protein solubility in a heterologous expression system.

## Materials and methods

### Evolutionary analysis by Maximum Likelihood method

Amino acid sequences were retrieved from The Arabidopsis Information Resource (TAIR; Berardini et al., 2015) for Arabidopsis sequences and the Marchantia polymorpha genome database (Marpol Base; Tanizawa et al., 2025). Sequences were aligned using the MUSCLE alignment method (Edgar 2004) in MEGA X (Kumar et al., 2018). Evolutionary analyses were conducted in MEGA X (Kumar et al., 2018).

### Plant growth conditions

*Marchantia* was grown under axenic conditions on a ½ Gamborg B5 solid media with 1% agar in a 90 mm X 25 mm petri dish. Plants were grown in a controlled growth room with a 16 h light and 8 h dark cycle with a light intensity of 120 mmol m^-2^ s^-1^ at 21 °C and constant relative humidity of 70%. The lid was sealed with 3M micropore surgical paper tape. To induce reproductive structures, the media was supplemented with 1% Glc and plants were grown under additional light in the far-red spectrum (720-740 nm).

### Agrobacterium tumefaciens transformation

30 µl of electrocompetent *A. tumefaciens* (strain GV3101) was inoculated with 1 µl of purified plasmid DNA. The electrocompetent cells were transformed with a 2 mm electroporation cuvette using the BIO-RAD Micropulser™. After electroporation, 700 µl of liquid 2X Yeast Extract Tryptone (2YT) medium (16g Tryptone, 10g yeast extract, 5g NaCl, in 1L ddH_2_O, pH 7.0) was added to the cuvette and mixed by pipetting. 500 µl of the culture was transferred to a 1.75 ml Eppendorf tube and shaken for 1 h at 28 °C with shaking at 180 rpm. 50 µl of the culture was spread on 2YT media plates with appropriate antibiotics and 50 µg/ml rifampicin and grown for 2 d at 28° C.

### *Agrobacterium*-mediated transformation of *Marchantia* sporelings

Sporeling transformation was performed according to Ishizaki et al., (2008). Briefly, at the clean bench, a tube of sterilised sporangia was resuspended with 100 µl of sterile ddH_2_O and crushed with a pipette tip. Then, 400 µl of water and 500 µl of sterilisation media were added and left to stand for 1 min. The spores were centrifuged for 1 min at 8000g, and the supernatant was removed. The spores were washed twice with 1 ml of sterile ddH_2_O and resuspended in 500 µl of sterile ddH_2_O. 100 µl of the spore suspension was added to 25 ml of Gamborg’s B5 liquid media (1.605g Gamborg B5 media, 2% sucrose, 0.03% L-glutamine, 0.1% casamino acids in 1L ddH_2_O, pH 5.5 with KOH) in a 125ml flask with a ventilated lid and grown for 7d on a shaker at 110 rpm. Two days before transfection, a single colony of *A. tumefaciens* was inoculated in 10 ml of 2YT media (16g Tryptone, 10g yeast extract, 5g NaCl, in 1 L ddH_2_O, pH 7.0) to grow for 2d in 30°C at 180 rpm. The 10 ml *A. tumefaciens* culture was centrifuged for 10 min at 3250 g. The supernatant was discarded, then resuspended in 10 ml of Gamborg’s B5 liquid media with 100 µM acetosyringone. The resuspended culture was incubated for 4 h at 30°C with shaking at 180 rpm. 1 ml of the *A. tumefaciens* culture was added to a 25 ml sporeling culture. For transformation using two constructs, 1 ml of each culture was added. The sporelings were co-cultured with *A. tumefaciens* shaking at 110 rpm under standard growth conditions for 2 d. The transfected sporelings were rinsed with 50 ml of ultrapure water over a 20-micron nylon mesh, then plated on appropriate antibiotic selection plates with timentin (300mg/L). After 2 weeks, the primary transformants were transferred to a second selection plate (T1 plants). A gemmae from a T1 plant was transferred to a third antibiotic selection plate to generate an isogenic transformant (T2 plant) for downstream analyses.

### Fertilisation of *Marchantia*

In a sterile bench, sterile ddH_2_O was pipetted onto the mature antheridiophore receptacles for 5 min to extract sperm from the antheridium. Droplets were drawn and pipetted onto the archegoniophore receptacles and left for 10 min. This process was repeated three times throughout a week.

### Subcellular localisation

Constructs used in this study are outlined in Table 3. The CD3-967 plasmid (Nelson et al., 2007) was used as a template to construct the Golgi organelle marker. CD3-967 uses the first 49 amino acids (aa) of the *Glycine max* (1,2)-α MANNOSIDASE I (GmMANI) protein to target the mCHERRY red fluorescent protein (RFP) to the Golgi (Saint-Jore-Dupas et al., 2006). The Golgi targeting sequence and mCHERRY was amplified from CD3-967 using CACC_CD3-967_Fwd and CD3-967_Rev (Table 4), cloned into the pENTR using the Gateway® pENTR-d-TOPO kit (Thermo Fisher Cat. No. K240020SP), then recombined into pMpGWB303 (Addgene # #68631) (Ishizaki et al. 2015) using the Gateway™ LR Clonase™ enzyme (Thermo Fisher, Cat. No. 11791-020). The translational reporter lines for MpARADL1 and MpARADL2 were constructed using overlapping PCR. 3.059 kilobases (kb) and 2.945kb upstream of the MpARADL1 and MpARADL2 start codons (promoter region) with a KpnI restriction site at the 5’ end, the coding sequences for MpARADL1 and 2 without the stop codon, and the GS linker-mNEON sequence (derived from pNEW) with a XbaI restriction side at the 3’ end were amplified with overlapping sequences between the components. In the overlapping PCR, the promoter region, coding sequence and GS-linker-mNEONgreen amplicons were added as template, and amplified in a single PCR reaction to generate the proARADL:ARADL-GS linker-mNEON amplicon. The amplicon was digested with KpnI and XbaI, and ligated into the pMIGRO entry vector, also digested with KpnI and XbaI using the T4 DNA ligase (New England Biolabs Cat No.1797 M0202S). The NotI cassette of pMIGRO was transferred to the binary vector pHART.

**Table 3.** Constructs used in this study.

| Vector | Construct |
| --- | --- |
| pHART | proMpARADL1(3.059kb):3XmVENUS |
| pHART | proMpARADL2(2.945kb):3XmVENUS |
| pHART | proMpARADL1:GUSplus |
| pHART | proMpARADL2:GUSplus |
| pHART | proMpARADL1:MpARADL1-mNEON |
| pHART | proMPARADL2:MpARADL2-mNEON |
| pMpGWB303 | proMpEF1:SmManI(1-49)-mCHERRY |
| pGE010 | proMpEF1:CAS9-proMpU6:MpARADL1gRNA1 |
| pMpGWB301 | proMpU6:MpARADL1gRNA2 |
| pGE010 | proMpEF1:CAS9-proMpU6:MpARADL2gRNA1 |
| pMpGWB301 | proMpU6:MpARADL2gRNA2 |
| pHART | proMpEF1:MpARADL1 |
| pHART | proMpEF1:MpARADL2 |
| pMpGWB303 | proMpEF1:amiR-MpARADL1+2<sup>Mpmir160</sup> |
| pKART | proMpARADL1:MpARADL1amiR-Res-mNEON |
| pKART | proMpARADL2:MpARADL2amiR-Res-mNEON |
| pDONR221 | TEV-MpARADL1(Δ1-36) |
| pDONR221 | TEV-MpARADL1(Δ1-44) |
| pDONR221 | TEV-MpARADL1(AA84-398) |
| pDONR221 | TEV-MpARADL1(Δ1-79) |
| pENTR | TEV-MpARADL2(Δ1-101) |
| pENTR | TEV-MpARADL2(AA159-476) |
| pDONR221 | TEV-MpARADL2(Δ1-151) |
| pDONR221 | TEV-AtARAD1(Δ1-33) |
| pDONR221 | TEV-AtARAD1(Δ1-37) |
| pENTR | TEV-AtARAD1(AA56-368) |
| pDONR221 | TEV-AtARAD1(Δ1-55) |

**Table 4.**
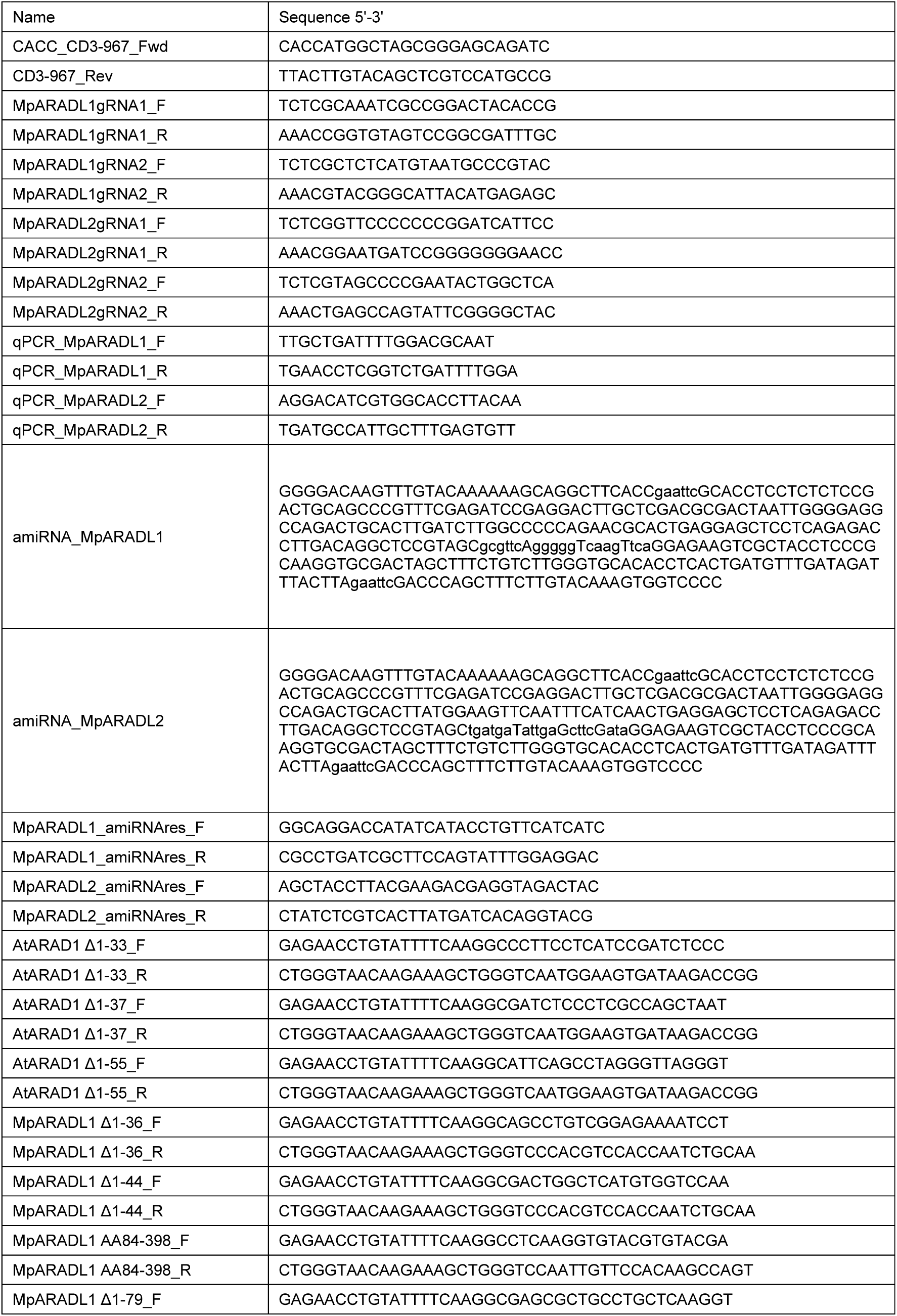
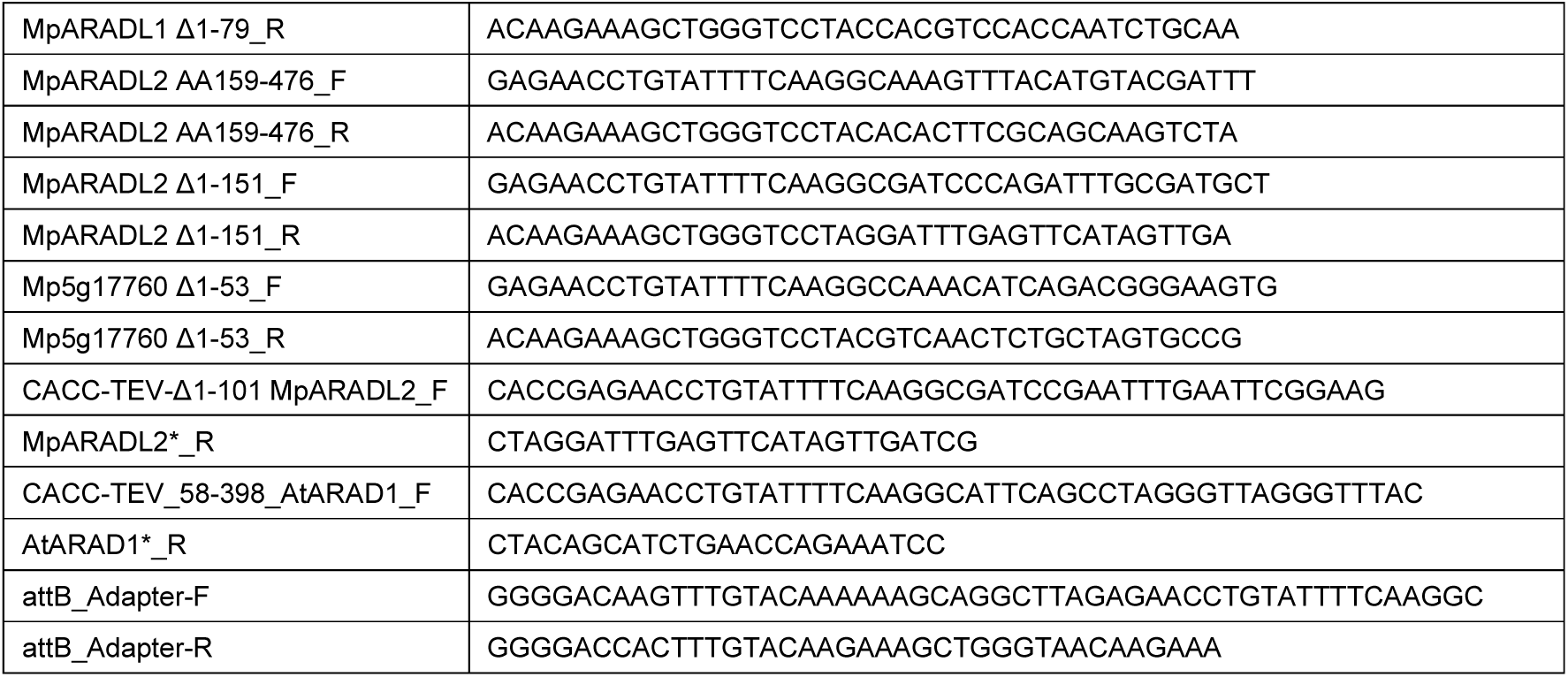
Primers used in this study.

### CRISPR

CRISPR-mediated gene editing was conducted according to Sugano et al. (2018). To build the CRISPR constructs, gRNA dimers were generated by incubating 5 µl of 10 uM forward and 5 µl of 10 uM reverse primers for each gRNA (Table 4) in the thermocycle at 95 °C for 5 min, 95°C to 85°C ramping at -2°C/s, 85°C to 25°C at -0.2°C/s and 1 min at 25°C. The dimerized gRNAs were ligated into a BsaI digested entry vector (MpU6-1_sgRNA_pENTR) using the T4 DNA ligase. Two gRNAs were designed for each gene. One gRNA was subcloned into pMpGE010 (Addgene #71536), and the second gRNA into pMpGWB301 (Addgene #68629) using the Gateway™ LR Clonase II. Sporelings were transformed as described above with the two plasmids simultaneously, and edits were confirmed through Sanger sequencing of the genomic DNA of transformants (AGRF, Australia).

### Overexpression

RNA was isolated from the thallus of wild-type Marchantia collected in Melbourne, Australia (RNeasy Mini Kit, Qiagen, Cat. No. 74104). 500ng of RNA was used for first-strand cDNA synthesis carried out using the Tetro™ Reverse Transcriptase (Meridian Bioscience, Cat no. BIO-65050). The coding sequences of *MpARADL1* and *2* with KpnI and BamHI restriction enzyme overhangs were amplified from the Marchantia cDNA and subcloned into a KpnI digested pBJ36 vector harbouring the Marchantia *Elongation Factor 1* (*MpEF1α*) promoter using the T4 DNA ligase. The NotI expression cassette of pBJ36 was subsequently transferred into the binary vector pHART.

Overexpression of MpARADL1 and 2 was confirmed at the transcript level using qRT-PCR. RNA from 2-week-old plants was extracted using the RNeasy Mini Kit. 500 ng of RNA was used for reverse transcription using Tetro™ Reverse Transcriptase. 4X diluted cDNA was used as a template for SYBR Green master mix (Bio-Rad, USA, cat no. #1725270). Three biological replicates were run in three technical replicates. Samples were analysed in Hard-Shell® 384-Well PCR Plates (BIO-RAD, USA, cat no. #HSP3805) using the Bio-Rad CFX384 Touch Real-Time PCR Detection System. The cycling condition was initial denaturation for 30 sec at 95°C, denaturation for 10 sec at 95°C, annealing for 20 sec at 60°C, repeating the denaturation and annealing 40 times, then melt curve analysis at 65°C to 95°C with 0.5°C/5 sec increments. Mp ADENINE PHOSPHORIBOSYL TRANSFERASE (MpAPT) and Mp ACTIN (MpACT) were used as reference genes (Saint-Marcoux et al. 2015). The primers qPCR_MpARADL1_Fwd and qPCR_MpARADL1_Rev was used to quantify MpARADL1 expression, and qPCR_MpARADL2_Fwd and qPCR_MpARADL2_Rev were used to quantify MpARADL2 expression. The efficiencies of primers for qRT-PCR obtained according to (Jozefczuk and Adjaye 2011) were 101%, 99%, 107% and 90% for MpACT, MpAPT, MpARADL1 and MpARADL2 respectively.

### amiRNA induced gene silencing

The amiRNA sequence targeting MpARADL1 and 2 were designed according to Flores-Sandoval et al., (2016). Briefly, MpARADL1 and 2 were aligned using CLUSTAL Omega (https://www.ebi.ac.uk/jdispatcher/msa/clustalo). From a region of high sequence similarity, a 21 bp sequence in the sense orientation, or the “target sequence” was chosen which ends with an adenine (A) and is GC rich in bases 3-5. This sequence was then reverse complemented to create the “amiRNA sequence”. Then, the duplex for the “amiRNA sequence” (mir*) was generated by introducing mismatches in the “target sequence” in positions 7 and 18 and creating G/U complementarity in position 13. With these sequences, oligos encoding the MpMIR160 microRNA hairpin sequence substituted with the “amiRNA sequence” and “mir*” sequence, flanked by Gateway® BP adapter sequence and an EcoRI restriction sites were ordered from Integrated DNA Technologies (https://eu.idtdna.com/pages), and cloned into the pDONR221 entry vector using Gateway® BP Clonase Enzyme II (Thermofisher, Cat. No. 11789013), then recombined into pMpGWB303using Gateway LR clonase II. Reduced expression of MpARADL1 and 2 was confirmed using qRT-PCR, as outlined above.

Constructs harbouring the amiRNA resistant copies of *MpARADL1* and *2* coding sequences were designed by introducing six silent mutations in the amiRNA target site. These coding sequences were generated by using the Q5 site directed mutagenesis kit (NEB: E0554S) following manufacturer’s protocols. Primers were designed using NEBaseChanger software (https://nebasechanger.neb.com/). As a base for the amiRNA resistant constructs. constructs for the translational reporter lines described above were used as template. Primers MpARADL1_amiRNAres_F and MpARADL1_amiRNAres_R and MpARADL2_amiRNAres_F and MpARADL2_amiRNAres_R were used to mutagenize the respective gene constructs (Table 4). In a 25 µl reaction, 12.5 µl of the Q5 Hot Start High-Fidelity 2X Master Mix, 1.25 µl of 10 µM forward primer, 1.25 µl of 10 µM reverse primer, 25 ng of template plasmid and up to 9 µl of nuclease free water was added. The mutagenized plasmid was subcloned into the pKART destination vector using the NotI restriction enzyme cassette.

### CoMPP

For each plant line, six technical replicates of 10 mg of de-starched AIR were made from 3-week-old thalli. AIR was generated according to (Rautengarten et al., 2019). The analysis was conducted according to Fradera-Soler et al., (2022).

### Glycosyl linkage analysis

Glycosyl linkage analysis was conducted using gas chromatography-mass spectrometry (GC-MS) of partially methylated alditol acetates (PMAAs) derivatised from AIR. AIR was generated from 3-week old plants, other than the amiRNA line, which required bulking over a period a several months. The analysis was performed as outlined in Zhang et al., (2024) and Black et al. (2023).

### Immunolabelling

Mature sporophytes were fixed for 1 h in 4% paraformaldehyde (Pro Sci Tech Cat. No. C006) and 0.5% glutaraldehyde (Sigma Cat. No. G5882-10X10ML) in phosphate buffered saline (PBS). Sporophytes were washed three times with PBS, 10 minutes each time then once with ddH_2_O for 10 min. To dehydrate the tissue, sporophytes were washed in 10%, 20%, 40%, 60%, 80% then 100% ethanol for 30 min each, then in another 100% ethanol wash overnight. Sporophytes were infiltrated with 50%, 75% LR white resin in ethanol, then in 100% resin three times for 8 h each. Resin infiltrated sporophytes were placed in BEEM capsules filled with LR white resin, then polymerised at 60 °C overnight. 1 µM sections were made using glass knives on the Leica Ultracut S Microtome. The resin sections were placed on superfrost Polysine coated microscope slides (Bio-Strategy, Cat. No. EPBRSF41296PL) with hydrophobic boundaries drawn with the Super Pap Pen (Sigma Aldrich Cat no. Z672548) and dried on a heat block until all the moisture was evaporated. Dried sections were incubated with a droplet of blocking buffer (PBS with 5% w/v skim milk powder) for 1 h at room temperature. Throughout the incubation process, the sections were protected from light. After each incubation, the droplets were removed by pipetting. After blocking, the sections were incubated with a primary antibody diluted 1:10 times in blocking buffer. Then, the sections were washed thrice for 5 min each with PBS. The sections were incubated with a secondary antibody (Goat anti-Mouse IgG (H+L) Highly Cross-Adsorbed Secondary Antibody, Alexa Fluor™ Plus 488, Cat No. A32723) diluted 1:1000 in the blocking buffer for 1 h at room temperature. The sections were washed thrice for 5 min each with PBS. Finally, the section was incubated with a droplet of 0.1M calcofluor white for 1 min and washed thrice for 5 min each with PBS. The sections were observed with the Leica DM6000 widefield compound microscope. Calcofluor white was excited at 380 nm and YFP was excited at 488 nm. Image acquisition was performed using the Metamorph Online Premier software, version 7.10.1.161.

### Scanning Electron Microscopy

Plant tissue for imaging was washed in 5%, 10%, 20%, 30% and 50% ethanol for 30 min each, then fixed in the fixing solution (50% ethanol, 5% acetic acid, 10% formaldehyde) for 1h. The samples were further washed in 60%, 70%, 80%, 90%, 100% ethanol for 30 min each. Samples were kept in fresh 100% ethanol overnight. Ethanol was exchanged from samples using the Leica EM CPD 300 critical point dryer, then sputter coated with gold using the Quorum QT5000 Sputter Coater. Samples were imaged using the Hitachi TM4000 II Tabletop SEM.

### HEK293 cell heterologous expression

For each protein of interest, the amino acid sequence was analysed with a tool to identify the transmembrane domain called Transmembrane Helices Hidden Markov Models v2.0 (TMHMM v2.0, Krogh et al. 2001, https://services.healthtech.dtu.dk/services/TMHMM-2.0/) which produces a probability score indicating whether an amino acid residue is likely to be ‘inside’ (cytosolic), ‘membrane’ (transmembrane) and/or ‘outside’ (luminal). Constructs were designed according to Urbanowicz et al. (2014). Truncated coding sequences of AtARAD1 were amplified from cDNA prepared from rosette leaves of *A. thaliana* (Col-0). Truncated coding region sequences of *MpARADL1* and *2* were amplified from Marchantia cDNA. For the first round of amplification, the TEV protease recognition sequence encoding Glu-Asn-Leu-Tyr-Phe-Gln-Gly, followed by the gene specific sequence was used as the forward primer (Table 4). For the reverse primers, a partial attB adapter sequence followed by the gene specific sequence. A second set of universal primers, attB_Adapter-F, and attB_Adapter-R, were used to amplify the PCR products from the first round of amplifications to complete the attB recombination region. The completed attB-TEV protease sequence-PCR products were gel purified using NucleoSpin Gel and PCR Clean-up XS (Macherey-Nagel, item no. 740611.250) and cloned into pDONR221 plasmid vector (Life Technologies) using Gateway® BP Clonase Enzyme II (Thermofisher, Cat. No. 11789013) following the manufacturer’s instructions. The transformants were confirmed by sequencing with the M13_ (-21) _F primer and M13_R. Two constructs, encoding Δ101MpARADL2 and AA56-368 AtARAD1 could not be amplified using the primers designed as described. Thus, these two constructs were amplified using primers CACC_TEV-Δ1-101_MpARADL2_F and MpARADL2*_R, and CACC_TEV_58 398_AtARAD1_F and AtARAD1*_R, respectively, then subcloned into pENTR entry vector using the Gateway® pENTR-d-TOPO kit. HE293 cell heterologous expression of the GT47B proteins was conducted according to Prabhakar et al. (2020).

